# Glucocorticoid receptor and RUNX transcription factors cooperatively drive CD8 T cell dysfunction in human cancer

**DOI:** 10.1101/2025.04.30.651472

**Authors:** Christopher J Ward, Soura Chakraborty, Sanu K Shaji, Clara Veiga-Villauriz, Aws Al-deka, Qiuchen Zhao, Jhuma Pramanik, Xi Chen, Bidesh Mahata

**Author notes:** Equal contribution (Christopher J Ward, Soura Chakraborty).

## Abstract

Glucocorticoids (GCs) are potent modulators of immune responses; however, the mechanisms by which GCs regulate gene expression in human CD8 T cells remain incompletely defined. Here, we delineate how physiological cortisol signalling shapes the transcriptional and chromatin landscapes of primary human CD8 T cells. We identify a substantial cohort of GC-responsive genes that are co-regulated through the cooperative activity of the glucocorticoid receptor (GR) and RUNX transcription factors. Integrative RNA sequencing and ChIP sequencing analyses identified genome-wide cortisol-responsive immunoregulatory genes. Genetic deletion of the GR (encoded by *NR3C1*) abolished cortisol-induced gene expression changes, confirming GR-dependency. Notably, GR chromatin occupancy in cortisol-treated CD8 T cells was strongly enriched at RUNX transcription factor (TF) motif rather than canonical GC response elements (GREs). Co-immunoprecipitation assays validated a ligand-dependent physical interaction between GR and RUNX and revealed the interacting domains. Single-cell transcriptomic analyses of tumour infiltrating CD8 T cells revealed significant enrichment of cortisol-responsive genes, indicating an active GC signalling (response) within the tumour microenvironment. GR-RUNX dual controlled genes were enriched in tumour-infiltrating CD8 T cells across multiple cancer types, including lung adenocarcinoma, head and neck squamous carcinoma, pancreatic cancer, and breast cancer. We found GR-RUNX co-regulated genes are predominantly expressed in the predysfunctional state of CD8 T cell population of different solid tumours. These results suggest that local cortisol signalling within tumour microenvironments drives CD8 T cell dysfunction through GR-RUNX TF cooperation. Collectively, our findings identify RUNX3 TF as a critical mediator of GR signalling in human CD8 T cells and reveal a novel mechanism by which endogenous GCs influence antitumour immunity, which could be therapeutically targetable.

## Introduction

Glucocorticoids (GCs) are steroid hormones widely recognised for their potent immunomodulatory and anti-inflammatory properties. Synthetic GCs, such as dexamethasone, betamethasone and prednisone, are extensively utilised in clinical settings to treat autoimmune disorders, inflammatory diseases, and haematological malignancies due to their ability to suppress immune cell activation and cytokine production^1,2^. Endogenous GCs, primarily cortisol in humans, also play critical roles in regulating immune homeostasis and responses under physiological and stress conditions^3^. Chronic or dysregulated GC signalling has been implicated in impaired antitumour immunity and cancer progression^2,4,5–10^. Despite extensive studies on synthetic GC-mediated immunosuppression, the precise molecular mechanisms by which physiological cortisol signalling modulates human CD8 T cell functions, particularly within tumour microenvironments, remain incompletely understood.

The GC receptor (GR), encoded by the *NR3C1* gene, is a ligand-activated transcription factor belonging to the nuclear receptor superfamily. Upon ligand binding, GR translocates into the nucleus, binds chromatin at GC response elements (GREs), and regulates target gene expression through direct DNA binding or tethering interactions with other transcription factors^11^. Although classical GRE-dependent mechanisms have been extensively characterised in various cell types, emerging evidence suggests that GR can also regulate gene expression through interactions with non-canonical transcriptional partners. GR can modulate gene expression in immune cells through protein–protein interactions with other transcription factors such as NF-κB and AP-1^12^. This “tethering” or “composite” model of gene regulation allows GR to repress or activate a distinct subset of genes indirectly, without binding DNA at traditional GRE motifs. For example, GR can bind to NF-κB p65 subunits, preventing NF-κB from inducing proinflammatory cytokine transcription. Conversely, it can cooperate with AP-1 to enhance the expression of anti-inflammatory mediators. Such interactions enable GR to modulate gene expression programs in a highly context-dependent manner^13,14^. Identification of these non-canonical GR partners in specific immune subsets is crucial for understanding the complex regulatory networks orchestrated by GCs in a physiological or pathological context of immune regulation.

CD8 T cells are central players in adaptive immunity, mediating cytotoxic responses against infected cells (such as virus infected) or cancer cells. Effective CD8 T cell responses are critical for tumour control and elimination. However, persistent antigen exposure and immunosuppressive signals within tumour microenvironments frequently drive CD8 T cell dysfunction or exhaustion, limiting their antitumour efficacy^15,16^. In the context of CD8 T cells, GCs have been shown to suppress T cell effector programmes, including the upregulation of checkpoint inhibitory proteins (CTLA-4, PD-1, LAG-3, TIM3) in a context dependent manner^5,17,18^. Recent studies indicate that the local GC signalling may contribute significantly to immune suppression within tumour microenvironment by promoting T cell dysfunction^2,4,5–7^, yet the underlying transcriptional mechanisms remain largely unexplored.

RUNX transcription factors represent a family of evolutionarily conserved proteins crucially involved in lineage specification, differentiation, proliferation, and functional regulation of various immune cell subsets^19^. RUNX proteins (all three paralogues RUNX1-3) bind DNA through conserved Runt domains (>95% sequence identity) at core consensus motif YGYGGTY to regulate gene expression programs essential for hematopoietic development and immune function. Among the RUNX family members (RUNX1-3), RUNX2 is predominantly known for its roles in osteogenesis. However, recent evidence reveals its broader involvement in immune regulation and cancer biology^20^. RUNX1 and RUNX3 are critical regulators of CD8 T cell development and function. During thymopoiesis, they collaborate to repress CD4 and enforce CD8 lineage commitment^21,22^. In mature CD8 cytotoxic T lymphocytes (CTLs), RUNX3 dominates. It directly binds to the enhancers (e.g., *Cd8a* enhancer E8I) to maintain CD8 expression^21,22^ and activating effector genes like *Ifng, Gzmb* (granzyme B), and *Prf1* (perforin) ^21,22^. Runx3 deficiency severely impairs cytolytic activity and proliferation^21,22^, while also promoting tissue-resident memory (T_RM_) CD8 T cells by upregulating tissue-retention genes and suppressing recirculation^23^. Runx1 supports thymocyte transitions and mature T cell homeostasis but plays a secondary role in peripheral CTLs^24^. Runx2, though largely restricted to early thymic development, contributes to CD8 memory cell maintenance^1,23^. Together, Runx3 emerges as the central driver of CD8 T cell identity, effector function, and tissue residency, while Runx1 and Runx2 provide developmental and memory-related support. The potential interaction between GR signalling and RUNX transcription factors in CD8 T cells has not been previously investigated.

In this study, we aimed to elucidate how physiological cortisol signalling modulates human CD8 T cell function at the molecular level. Using integrative RNA sequencing (RNA-seq) and ChIP sequencing (ChIP-seq) approaches combined with genetic deletion and biochemical validations, we demonstrate that cortisol profoundly reshapes the transcriptional landscape of primary human CD8 T cells through ligand-dependent GR-RUNX cooperation. We further provide single-cell transcriptomic evidence that this novel GR-RUNX regulatory axis operates within tumour-infiltrating CD8 T cells across multiple cancer types. Our findings identify RUNX as a critical non-canonical partner of GR in human CD8 T cells and suggest a previously unrecognised mechanism by which local endogenous GC signalling within tumour influence antitumour immunity, offering new therapeutic opportunities aimed at enhancing cancer immunotherapy efficacy by targeting context-specific GR interactions.

## Results

### Cortisol reshapes the transcriptional landscape and signalling pathways in human activated CD8 T cells

To investigate how physiological cortisol influences the transcriptional profile of human CD8 T cells, we performed RNA-seq on *in vitro* activated primary CD8 T cells treated with cortisol (100 nM, 48 hours) or vehicle control (Figure 1a). Differential gene expression analysis identified 297 significantly regulated genes (169 upregulated, 128 downregulated) (Figure 1b). Among the most strongly induced genes were *TSC22D3, IL7R, FKBP5 and PIK3IP1*. Notably, *PIK3IP1,* a negative regulator of PI3Ks, *and* amphiregulin (*AREG*), a cytokine implicated in tissue repair and immunosuppression, emerged as a novel cortisol-induced gene in CD8 T cells. A complete list of these differentially expressed genes can be accessed in the Supplementary Table S1. Gene set enrichment analysis (GSEA) using the Hallmark gene set collection revealed distinct patterns of pathway regulation following GC treatment (Figure 1c, Supplementary Table S2). Cortisol treatment enriched gene sets associated with cell cycle control and metabolic regulation including E2F targets, MYC targets, G2M checkpoint, and mTORC1 signalling. Conversely, inflammatory and effector-associated programmes were significantly supressed including inflammatory response, TNFα signalling via NF-κB, IL6/JAK/STAT3 signalling, apoptosis, and IL2/STAT5 signalling. These data suggest cortisol enforces a metabolic checkpoint state in CD8 T cells, limiting inflammatory effector functions while simultaneously promoting cell survival and proliferation-associated pathways.

**Figure 1.**
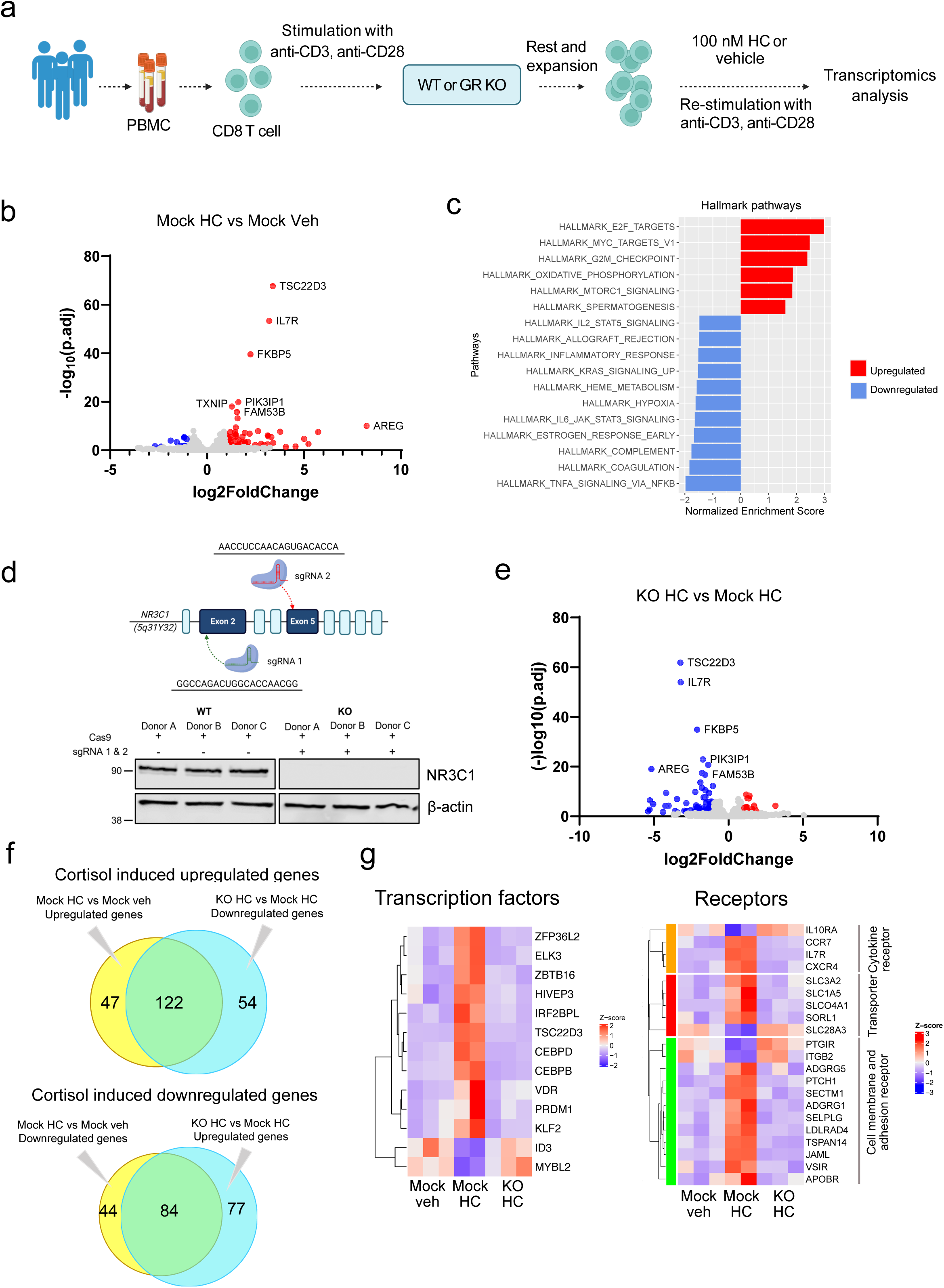
Genome-wide effect of glucocorticoid signalling in activated CD8 T cell. **a.** Schematic flowchart of the experimental strategy used to identify glucocorticoid-regulated genes in human CD8 T cells. CD8 T cells from three healthy donors were purified, activated, and expanded. *NR3C1* (encoding glucocorticoid receptor) was deleted in half of the cells. Wild type and GR-knockout CD8 T cells were then reactivated in the presence or absence of physiological levels (100 nM) of cortisol for 48 hours, to comprehensively identify glucocorticoid-dependent gene expression changes in CD8 T cells by RNA-seq. Specification of the samples (b-g): KO denotes GR knockout, Mock denotes wildtype, HC denotes hydrocortisone (cortisol) treated. Veh denotes vehicle. **b.** Volcano plot depicting the differentially expressed genes in human primary activated CD8 T cells after cortisol treatment. The significantly upregulated genes have been shown in red and the significantly downregulated genes have been shown in blue. Log2-Fold change versus –log10(adjusted p-value) has been plotted, n = 3 human samples in each group. **c.** The hallmark biological processes associated with cortisol-treated CD8 T cells. Prognosis-ranked gene set enrichment analysis (GSEA) identified hallmark pathways that are differentially regulated. Normalised enrichment scores (NES) were calculated for all 50 hallmark pathways in GSEA. Upregulated (shown in red) and downregulated (shown in blue) gene sets are displayed, with their NES values represented on the x-axis. **d.** Knockout (KO) of glucocorticoid receptor encoding gene *NR3C1* in CD8 T cells using CRISPR-Cas9 targeting exon 2 and exon 5 (top panel). The knockout efficiency has been validated by western blot (bottom panel). **e.** Volcano plot showing the differentially expressed genes in *NR3C1* KO activated CD8 T cells compared to mock activated CD8 T cells, in the presence of cortisol. **f.** The intersections in Venn diagrams showing the cortisol-induced GR-dependent upregulated and downregulated genes. **g.** Expression heatmap showing the cortisol-induced transcription factors and receptors (transcription factor, cytokine receptor, transporters, cell surface and adhesion receptor) from differentially expressed genes (log2FC> 1, log2FC <-1 and p.adj < 0.05).

To confirm GR-dependency of these transcriptional changes, we employed CRISPR-Cas9-mediated knockout of the GR (*NR3C1*) in primary human CD8 T cells (Fig 1d). Loss of GR expression abolished cortisol-induced upregulation of target genes (e.g., *TSC22D3*, *IL7R, FKBP5, PIK3IP1 and AREG*), confirming direct GR involvement in mediating cortisol-driven transcriptional reprogramming (Figure 1e). A complete list of differentially expressed genes in *NR3C1* knocked out CD8 T cell can be found in Supplementary Table S1.

In CD8 T cells, these 122 genes get upregulated in hydrocortisone (cortisol) treatment but downregulated when the GR (*NR3C1*) is knocked out. The Venn diagram analysis demonstrated that 47 genes were exclusively upregulated by cortisol in a GR-independent manner (Mock HC vs. Vehicle), while 54 genes were uniquely upregulated in GR-KO cells independent of the ligand cortisol (Figure 1f). 122 genes were upregulated in both conditions, suggesting GR-dependent mechanisms of cortisol-induced gene activation. Similarly, we found 84 genes get downregulated in cortisol treated CD8 T cell and get upregulated in *NR3C1* KO CD8 T cells. These findings identify cortisol induced true upregulated and downregulated genes in CD8 T cells. These findings also indicate that while a substantial proportion of cortisol-regulated genes require functional GR signalling, a subset of genes is expressed in a cortisol or GR-independent mechanisms.

To gain further insight into the functional implications of all cortisol-responsive genes (including both GR-dependent and GR-nondependent genes), we examined differentially expressed transcription factors and various receptors (Figure 1g). The heatmap analysis revealed distinct expression patterns of transcription factors regulated by cortisol treatment in both GR-sufficient and GR-deficient conditions. Several transcription factors, including *ZBTB16, ELK3, TSC22D3, VDR and CEBPB*, showed pronounced regulation following cortisol treatment. The receptor expression analysis identified multiple classes of differentially expressed receptors, including cytokine receptors (e.g., *IL10RA, CCR7, IL7R, CXCR4*), transporters (e.g., *SLC3A2, SLC1A5*), cell surface and adhesion receptors (e.g., *PTGIR, ITGB2, ADGRG5*). Among these, several interleukin receptors showed predominantly GR-dependent regulation, while others exhibited mixed patterns of GR-dependent and independent expression changes. These findings highlight the complex interplay between GR-dependent and independent mechanisms in mediating the effects of cortisol on the transcriptional landscape of CD8 T cells, with potential implications for understanding GC actions on immune function.

Together, these results demonstrate that physiological cortisol reshapes the transcriptional landscape of human CD8 T cells through GR-dependent regulation of immunomodulatory genes and pathways, highlighting a novel role for GCs in controlling T cell functionality via metabolic checkpoint enforcement.

### Genome-wide chromatin occupancy reveals the direct target genes of GR and its noncanonical association with RUNX transcription factors

To elucidate the molecular mechanisms underlying cortisol-mediated transcriptional reprogramming in activated CD8 T cells, we performed chromatin immunoprecipitation followed by sequencing (ChIP-seq) to map genome-wide GR binding sites upon cortisol treatment. Primary human CD8 T cells from healthy donors were activated *in vitro* and treated with cortisol (100 nM, 48 hours) or vehicle (Figure 2a). GR-bound chromatin regions were identified by ChIP-seq. Cortisol elicited a robust GR chromatin-binding response, yielding 4,871 high-confidence GR peaks relative to vehicle controls (Figure 2b; Supplementary Table S3). Motif enrichment analysis of these GR-bound genomic regions surprisingly revealed no significant enrichment of canonical GREs, however, showed a striking enrichment of non-canonical enrichment of RUNX transcription factor binding motifs upon cortisol treatment (Figure 2b). Specifically, RUNX motifs were present in approximately 12.6% (Donor A) and 35.92% (Donor B) of GR-bound peaks in cortisol-treated cells. Other transcription factor motifs such as TEAD and NFAT also showed modest enrichment following cortisol stimulation (Figure 2b). These data suggest that cortisol-induced GR chromatin occupancy in CD8 T cells preferentially occurs at genomic loci harbouring non-canonical binding to other TFs sites, particularly RUNX binding sites rather than canonical GREs. However, we observed donor-to-donor variability. Cross-donor comparison of Donor A and Donor B ChIP-seq analysis of CD8 T cell identifies 1103 common peaks between donors CD8 T cells treated with hydrocortisone (cortisol) (Figure 2c). Despite the donor-to-donor variability, 344 overlapped peaks were detected between vehicle-treated and hydrocortisone (cortisol) treated samples (Figure 2d). It indicates extensive cortisol-dependent remodelling of GR occupancy in CD8 T cells. We identified 81 putative direct targets of GC by integrating GR-occupied genomic regions from ChIP-seq and cortisol-induced upregulated genes (Figure 2e). We also observed, 50 of these 81 target genes were associated with GR peaks containing RUNX motif, revealed their coregulation by GR-RUNX interaction. (Figure 2f). Notably, this gene set included critical immunoregulatory targets induced by cortisol such as *TSC22D3, IL7R, FKBP5 and PIK3IP1* (Figure 2g). Further genomic annotation analysis indicated that cortisol-induced GR binding predominantly localised to intronic (31%) and promoter-proximal regions (31%), supporting a regulatory network distinct from classical GRE driven enhancer occupancy. (Supplementary figure S1a). Collectively, these data implicate RUNX-associated chromatin engagement as dominant feature of GR-dependent gene regulation in activated CD8 T cell. The list of GR direct target genes can be found in Supplementary Table S4.

**Figure 2.**
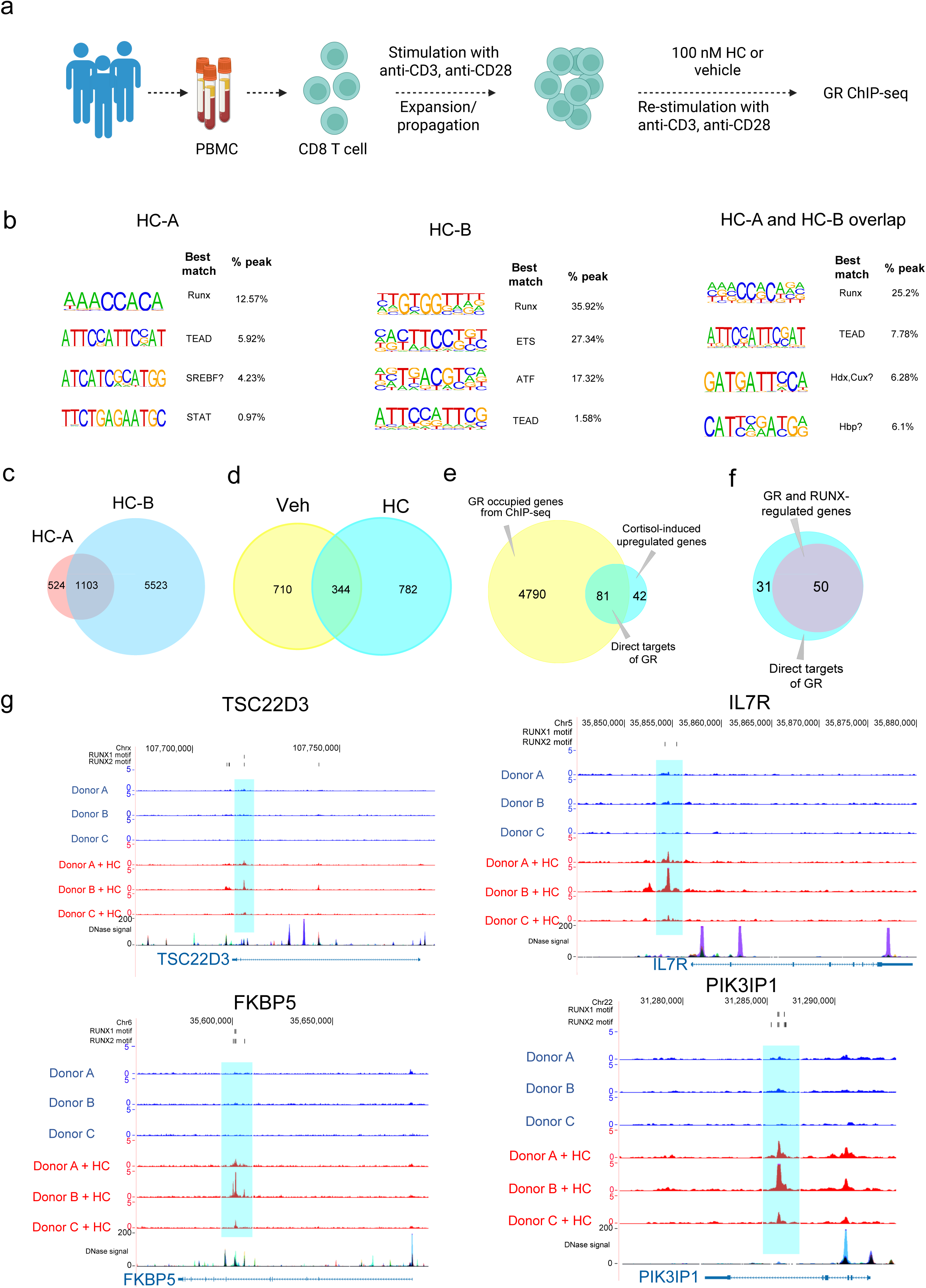
Genome-wide chromatin binding of GR identifies non-canonical transcription factor binding motif in CD8 T cells. **a.** Diagrammatic flowchart of the experimental strategy. CD8 T cells from healthy donors were purified, activated and propagated. Resting CD8 T cells were reactivated the presence or absence of physiological level (100 nM) cortisol to find out genome-wide chromatin binding of GR by ChIP-seq. **b.** Homer motif analysis of cortisol-treated CD8 T cells. The top 4 enriched GR binding peaks and corresponding motifs have been shown for two healthy donors (HC-A & HC-B) in cortisol-treated CD8 T Cells. The motifs have been arranged in descending order of abundance/enrichment. For each motif, the best match transcription factor and percentage of GR peaks containing the motif are indicated. HC-A & HC-B: Hydrocortisone (HC)/cortisol-treated donor-A & donor-B samples. **c.** Venn diagram showing the common number of peaks between hydrocortisone (cortisol) treated CD8 T cells from two donors. **d.** Venn diagram showing glucocorticoid (GC)-induced peaks comparing vehicle-treated and cortisol (HC)-treated CD8 T cells. **e.** The Venn diagram shows the direct targets of glucocorticoids after intersecting GR binding peaks from ChIP-seq data and cortisol-induced upregulated genes. **f.** The Venn diagram shows all GR-RUNX co-regulated genes that are also direct targets of GR. **g.** Genome browser tracks showing GR ChIP-seq signal in activated CD8 T cells from three independent healthy donors (donor A-C) treated with vehicle or hydrocortisone (HC). The glucocorticoid-influenced and GR-RUNX-regulated four genes are shown here. The shaded region highlights the cortisol-induced GR peaks with the predicted RUNX motif annotated above the peaks. Dnase I hypersensitivity signal is shown as an indicator of chromatin accessibility.

### Ligand-dependent interaction between GR and RUNX3 orchestrates cortisol-induced gene regulation in CD8 T cells

Given the prominent enrichment of RUNX motifs at GR-bound genomic regions (Figure 2), we hypothesised that GR might physically interact with RUNX transcription factors in a ligand-dependent manner. To test this hypothesis, first we compared the expression level of RUNX transcription factor in human CD8 T cells and found RUNX3 is predominantly expressed (Supplementary figure S1b). We therefore tested whether GR can physically interact with RUNX3 upon GC stimulation. We performed co-immunoprecipitation (co-IP) experiments in HEK293T cells transiently expressing GR and RUNX3 in the presence or absence of cortisol stimulation (100 nM, 36 hours) (Figure 3a). To define interacting domains of RUNX3 and GR for this GR-RUNX3 interaction, based on the previous research, we deleted specific domains of GR (NR3C1) and RUNX3 (Figure 3b)^25,26^. We observed GR and RUNX3 physically interact (Figure 3c). Removal of the GR N-terminal AF1 transactivation region (ΔAF1-NR3C1) substantially diminished RUNX co-precipitation compared with WT GR, indicating that the AF1-containing N-terminus contributes critically to stable GR–RUNX complex formation (Figure 3c). Conversely, deletion of the RUNX3 transactivation and inhibitory regions (ΔTAD ID-RUNX3) also weakened the interaction between RUNX3 and GR, demonstrating that RUNX3 C-terminal regulatory domains are required for efficient association with GR (Figure 3c). ΔTAD RUNX3 deletion mutation was not enough to diminish this interaction (Figure 3c). This co-IP analysis also demonstrated that RUNX3 co-precipitated with GR only upon cortisol treatment, confirming ligand-dependent formation of a GR-RUNX3 protein complex (Figure 3c).

**Figure 3.**
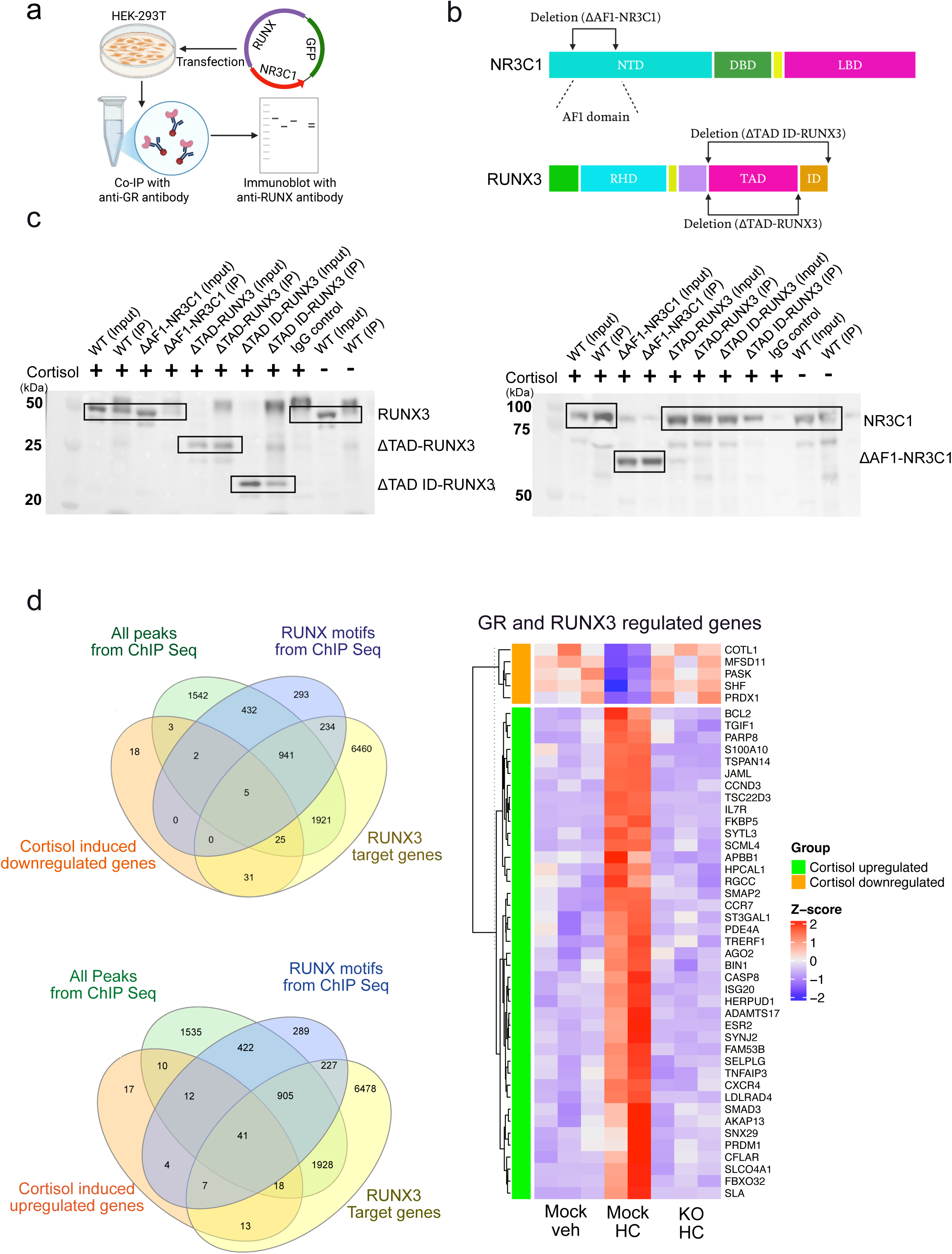
RUNX3 transcription factor cooperate with the glucocorticoid receptor to regulate gene expression. **a.** Schematic illustration of the experimental strategy (left). HEK 293T cells were transfected with GR and RUNX3 expressing plasmid construct. Cell lysates were immunoprecipitated by using anti-GR antibody and immunoblotted by RUNX antibody. Representative immunoblot shown in the right panel. **b.** Domain architecture of GR (NR3C1) and RUNX3, and the deletions used to map the interaction interface. For GR, the N-terminal AF-1 (NTD) region was deleted (ΔAF1-NR3C1). For RUNX3, the C-terminal transactivation/inhibition region was deleted (ΔTAD-RUNX3 and ΔTAD-ID-RUNX3). **c.** Co-IP mapping of the GR–RUNX3 interaction interface. Anti-GR (NR3C1) co-immunoprecipitation was performed in HEK-293T cells expressing wild-type (WT) GR and RUNX3 or the indicated deletion mutants (ΔAF1-NR3C1, ΔTAD-RUNX3, ΔTAD/ID-RUNX3). To assess ligand dependence, HEK-293T cells co-expressing WT GR and RUNX3 were treated with cortisol (+) or vehicle (–) prior to anti-GR co-IP. Input and immunoprecipitated fractions were immunoblotted with anti-RUNX3 and anti-GR (NR3C1) antibodies. **d.** Venn diagrams showing the GR-RUNX3 co-regulated genes by integrating cortisol induced genes (upregulated and downregulated), all GR binding peaks from ChIP-seq, GR binding site containing RUNX motif and RUNX3 target genes. The corresponding heatmap showing the expression of GR-RUNX3 co-regulated genes in vehicle-treated CD8 T cells, cortisol (hydrocortisone or HC) treated CD8 T cell and *NR3C1* KO CD8 T cells.

To assess the functional relevance of this interaction in primary CD8 T cells, we integrated our GR ChIP-seq, RNA-seq with previously publicly available RUNX target genes datasets that were reported based on RUNX ChIP-seq and RNA-seq. Integrating cortisol-responsive genes containing proximal GR peaks enriched for RUNX motifs with RUNX1 target genes identified 12 genes that showed strong cortisol induction together with nearby GR–RUNX motif occupancy, consistent with direct co-regulation by GR and RUNX transcription factors (Supplementary figure S1c). Next, we performed analogous integration with RUNX2 target genes and stratified the genes that are upregulated (45 genes) or downregulated (7 genes) by the cortisol induced GR-RUNX complex (Supplementary figure S1d). Using RUNX3 focused gene set integration in CD8 T cell, we observed 41 genes co-upregulated by GR-RUNX3 and 5 genes co-downregulated by GR-RUNX3 axis (Figure 3d). All the gene names corresponding to this figure have been listed in supplementary Table S5.

Collectively, these results demonstrate a model in which cortisol promotes physical interaction and functionally transcriptional cooperation between GR and RUNX3 in CD8 T cell. This ligand-dependent GR-RUNX interaction expands the regulatory repertoire of GCs beyond classical GRE-mediated transcriptional control, highlighting a previously unrecognised molecular mechanism by which endogenous GCs modulate human CD8 T cell function.

### GR-dependent cortisol-induced gene signature is enriched in tumour-infiltrating CD8 T cells across multiple cancer types

Given our findings that cortisol signalling profoundly reshapes the transcriptional landscape of human CD8 T cells via GR-RUNX cooperation, we next explored whether this regulatory axis is relevant in the tumour microenvironment. To address this, we reanalysed publicly available single-cell scRNA-seq datasets from tumour-infiltrating lymphocytes derived from patients with breast cancer (Figure 4a), lung adenocarcinoma (Figure 4b), head and neck squamous cell carcinoma (HNSCC) (Figure 4c) and pancreatic adenocarcinoma (Figure 4d). We scored steroid responsiveness in tumour infiltrating CD8 T cell and matched blood derived or matched tissue resident CD8 T cells based on the cortisol-responsive gene signature identified in our primary human CD8 T cell model (including key genes such as *TSC22D3, IL7R, FKBP5, PIK3IP1* and *AREG*. Strikingly, single-cell transcriptomic analyses revealed a consistent and significant enrichment of cortisol-induced genes in tumour-infiltrating CD8 T cells across all four cancer types examined (Figure 4a-d), suggesting potential heightened responsiveness to local GC cues.

**Figure 4.**
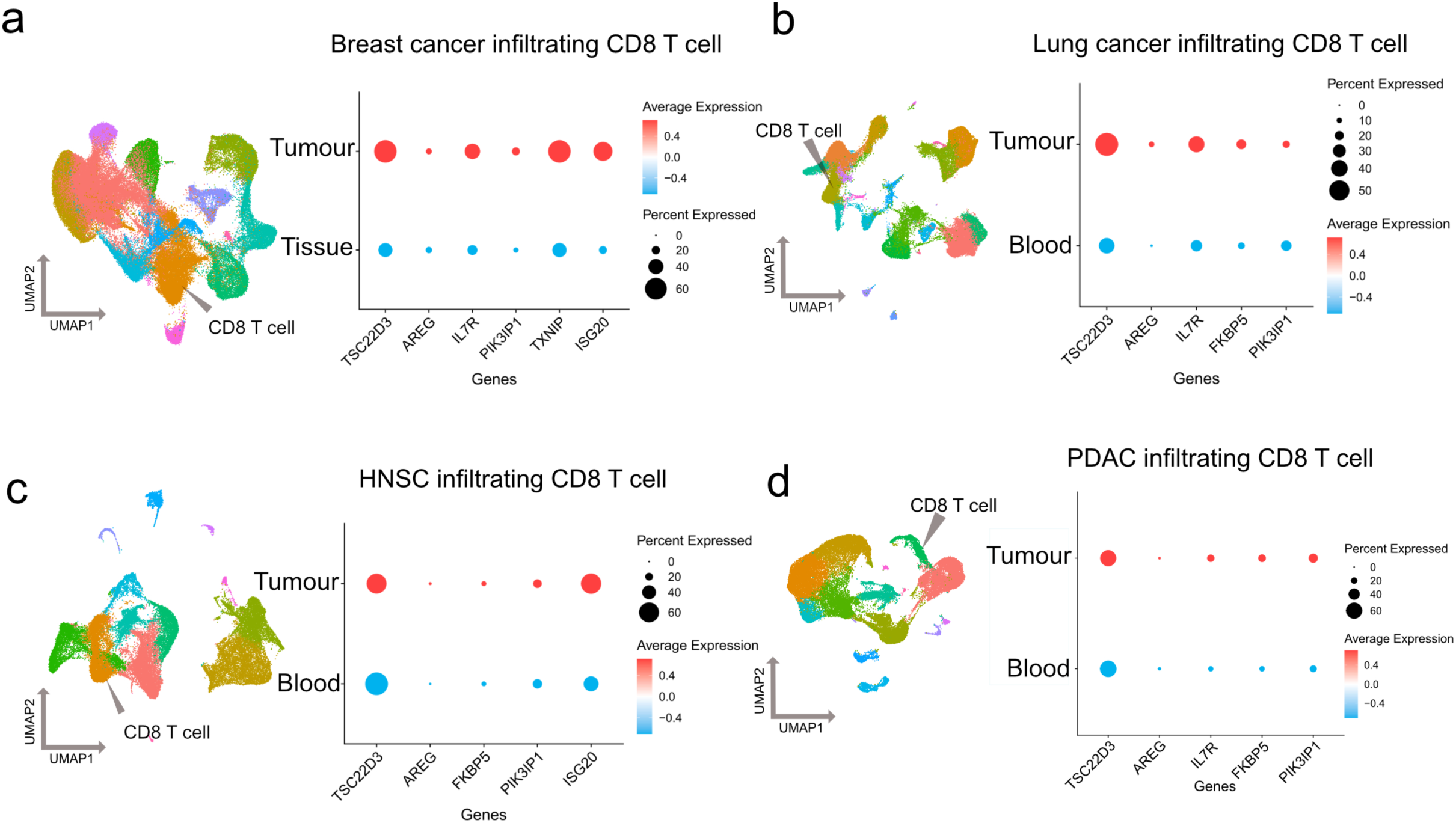
Single-cell transcriptomics study reveals the expression of cortisol responder genes in different tumour-infiltrating CD8 T cells. **a-d.** UMAP-based clustering of scRNA-seq data was used to identify CD8 T cell populations from tumour and matched blood or normal tissue samples in breast cancer (**a**), lung cancer (**b**), head and neck squamous cell carcinoma (HNSC) (**c**), and pancreatic ductal adenocarcinoma (PDAC) (**d**). Dot plots show the proportion of cells (dot size) and the average expression (colour intensity) of selected top cortisol responder genes within tumour-infiltrating CD8 T cells and corresponding peripheral blood or matched tissue-resident CD8 T cells. The data highlight distinct patterns of steroid-responsive gene expression between tumour-infiltrating and non-tumour CD8 T cell populations, reflecting the influence of the tumour microenvironment on CD8 T cell transcriptional states and potential links to T cell dysfunction and immune evasion

These observations indicate that endogenous GC signalling actively operates within tumour microenvironments to modulate CD8 T cell function, potentially contributing to an immunosuppressive state that favours tumour immune evasion. Collectively, these findings underscore the clinical relevance of GR-mediated transcriptional regulation in shaping immune responses within tumours and highlight the potential therapeutic value of targeting GR to reinvigorate antitumour immunity.

### Elevated expression of RUNX-GR co-regulated genes in tumour-infiltrating CD8 T cells across cancer types

We next investigated the expression patterns and functional significance of RUNX-GR co-regulated genes in CD8 T cells across multiple cancer types. We derived a unified transcriptional readout of GR–RUNX cooperation. Specifically, we integrated GR–RUNX1-, GR–RUNX2- and GR–RUNX3-mediated upregulated gene sets (12, 45 and 41 genes, respectively from Supplementary figure S1, Figure 3d) and identified a shared 50-gene core signature. Publicly available single-cell transcriptomic datasets of CD8 T cells isolated from tumours and matched adjacent normal tissue or blood across four major malignancies: triple-negative breast cancer (TNBC), lung cancer, head and neck squamous cell carcinoma (HNSC), and pancreatic ductal adenocarcinoma (PDAC) were reanalysed (Figure 5a-d). Across all datasets, RUNX-GR co-regulated genes exhibited significantly elevated expression in tumour-infiltrating CD8 T cells compared to their non-tumour counterparts (Figure 5a-d). The dot plots both the fraction of the cells expressing each gene (dot size) and average expression level (colour intensity) were consistently increased in tumour infiltrating CD8 T cell population in these cancer types indicating broad activation of RUNX-GR programme within the tumour microenvironment.

**Figure 5.**
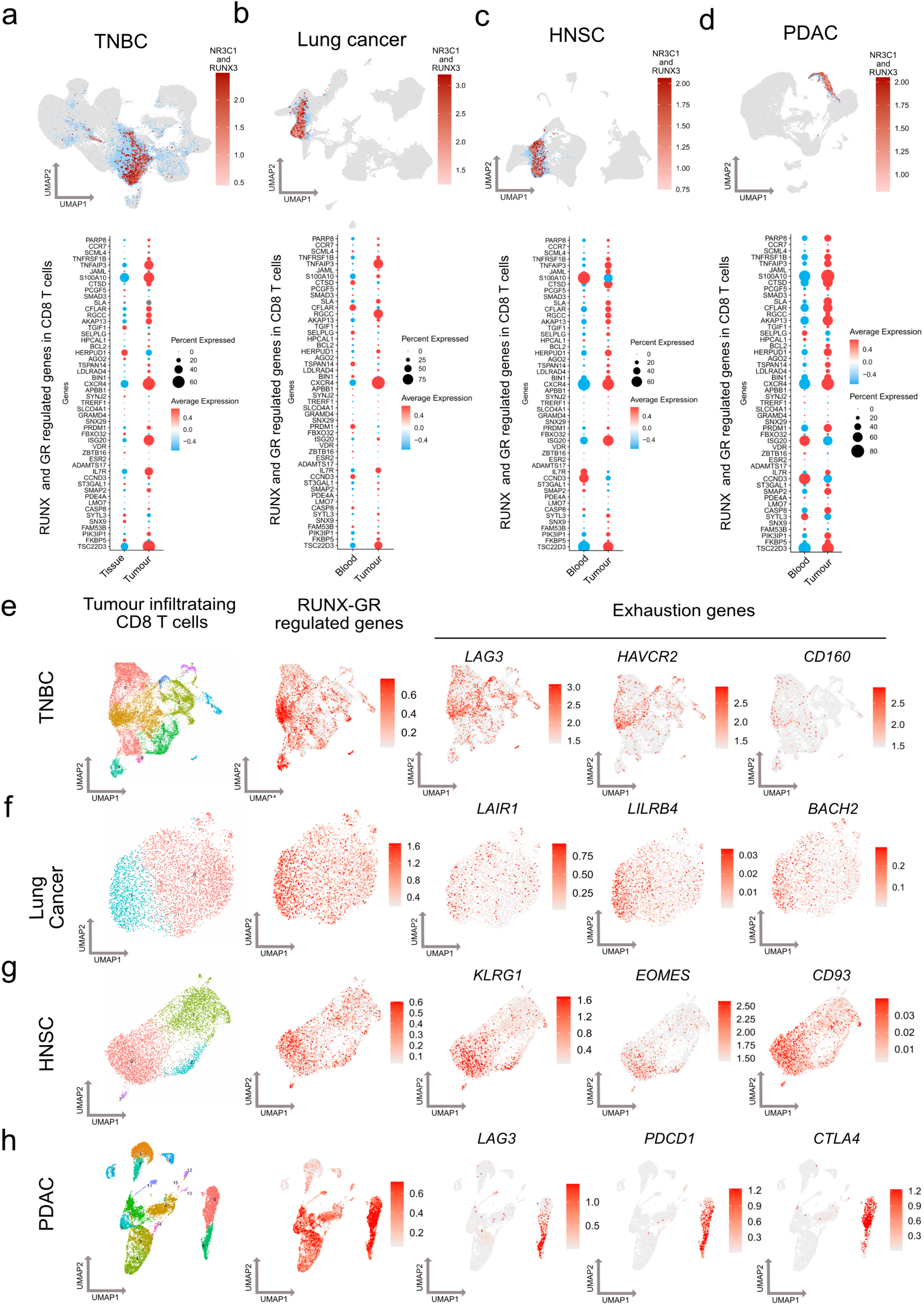
RUNX and GR co-regulated gene expression in tumour-infiltrating CD8 T cells across cancer types. **a–d,** Single-cell transcriptomic profiling of CD8 T cells isolated from tumours and matched non-tumour compartments (matched blood or matched non-cancerous tissue) in triple-negative breast cancer (TNBC) (a), lung cancer (b), head and neck squamous cell carcinoma (HNSC) (c), and pancreatic ductal adenocarcinoma (PDAC) (d). Top, UMAP feature plots showing the distribution of CD8 T cells (blue) with *NR3C1* (GR) and *RUNX3* expression (colour scale; grey indicates low/absent signal, red indicates higher expression). Bottom, dot plots summarising expression of the RUNX–GR co-regulated gene set across tumour-infiltrating CD8 T cells and matched non-tumour CD8 T cells (TNBC: adjacent tissue; lung/HNSC/PDAC: blood). Dot plots show expression of the RUNX–GR co-regulated gene set (a 50-gene core signature defined by integrating GR–RUNX1-, GR–RUNX2- and GR–RUNX3-mediated upregulated gene sets. Dot size indicates the fraction of cells expressing each gene and colour intensity denotes average expression. Across all tumour types, RUNX–GR co-regulated genes are expressed at higher levels in tumour-infiltrating CD8 T cells compared with their non-tumour counterparts . **e-h.** To assess the functional relevance of RUNX and GR co-regulated gene expression, module scores were calculated using the AddModuleScore method for each CD8 T cell. High module scores, indicative of elevated co-regulated gene expression, were associated with increased expression of exhaustion-related genes, highlighting a link between RUNX2/GR transcriptional activity and the exhausted phenotype in tumour-infiltrating CD8 T cells across TNBC **(e)**, lung cancer **(f)**, HNSC **(g)**, and PDAC **(h)**.

To assess the functional relevance of this expression pattern, we calculated per-cell module scores using AddModuleScore method (Figure 5e-h). Higher module scores, indicative of elevated RUNX-GR co-regulated gene expression, strongly correlated with increased expression of established T cell exhaustion/dysfunction markers. This correlation was consistent across TNBC (Figure 5e), lung cancer (Figure 5f), HNSC (Figure 5g) and PDAC (Figure 5h), highlighting a conserved link between RUNX-GR transcriptional activity and the exhausted phenotype in tumour-infiltrating CD8 T cells. These findings indicate that RUNX-GR dependent gene regulation is particularly prominent within the tumour microenvironment, suggesting its potential role in shaping the tumour-infiltrating CD8 T cells dysfunction In reanalysed single-cell RNA-seq datasets of tumour-infiltrating CD8 T cells from TNBC, lung cancer, HNSC and PDAC, we classified CD8 T cells into naïve-like, cytotoxic, pre-dysfunctional and dysfunctional states using established marker-gene signatures^27^. When the GR–RUNX co-regulated gene module was overlaid onto the same UMAP embeddings, cytotoxic CD8 T cells consistently formed a spatially distinct cluster from cells with high GR–RUNX module expression. Instead, GR–RUNX target-gene expression preferentially localised to regions enriched for the pre-dysfunctional programme and was partially adjacent to dysfunctional states, consistent with an association between GR–RUNX cooperation and acquisition of dysfunction in tumour-infiltrating CD8 T cells (Figure 6a-d).

**Figure 6.**
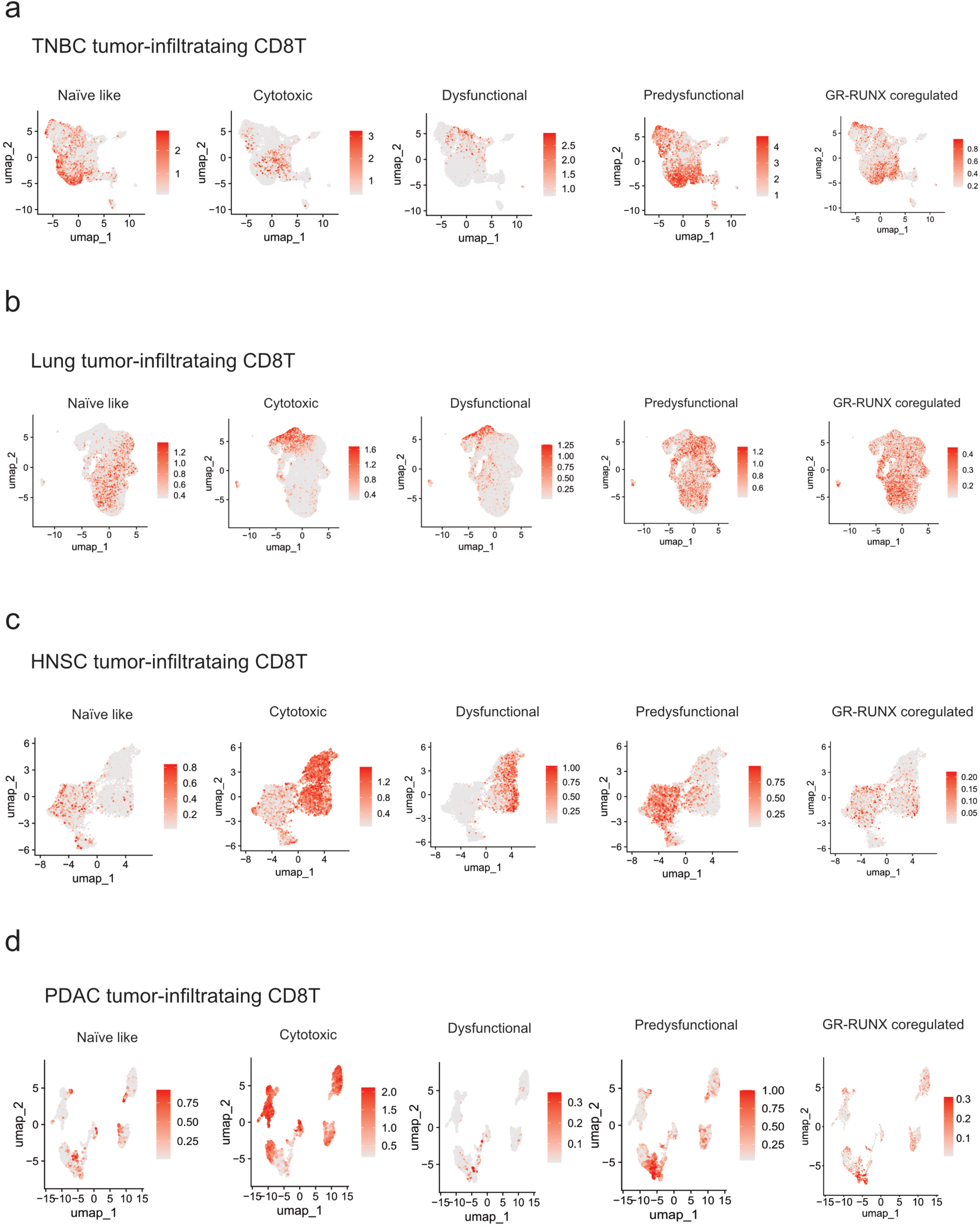
GR–RUNX co-regulated gene expression is enriched in pre-dysfunctional tumour-infiltrating CD8 T cells across solid cancers. **a–d,** Feature plots showing module scores for naïve-like, cytotoxic, dysfunctional and pre-dysfunctional CD8 T cell transcriptional states, together with the GR–RUNX co-regulated gene module in tumour-infiltrating CD8 T cells from **a,** triple-negative breast cancer (TNBC), **b,** lung cancer, **c,** head and neck squamous cell carcinoma (HNSC) and **d,** pancreatic ductal adenocarcinoma (PDAC). Colour intensity indicates relative module score (higher scores shown in darker red). Across tumour types, cytotoxic CD8 T cells make distinct cluster from GR–RUNX module–expressing CD8 T cells, whereas GR–RUNX module activity preferentially localises to areas enriched for the pre-dysfunctional programme and adjacent to dysfunctional states.

### Single-cell trajectory analysis reveals an association between cortisol-responsive gene expression and CD8 T cell dysfunction in cancer

To investigate how cortisol signalling affects CD8 T cell differentiation within the tumour microenvironment, we performed single-cell trajectory inference analysis on CD8 T cells isolated from lung tumours and paired peripheral blood samples from patients with lung cancer. Using the Monocle algorithm, pseudo-temporal trajectories were constructed based on single-cell transcriptomic profiles (Supplementary figure S2a). Cells expressing high levels of cortisol-responsive genes (*TSC22D3, IL7R, FKBP5*) were predominantly positioned along trajectory branches associated with progression towards terminal exhaustion or dysfunction, characterised by increased expression of inhibitory receptors, such as *PDCD1, CTLA4, HAVCR2, TIGIT, LAYN, and TOX* (Supplementary figure S2b). In contrast, cells exhibiting lower cortisol-responsive gene signatures were primarily associated with early activation states. These results suggest that GC signalling within the tumour microenvironment is activated shortly after CD8 T cells infiltrate the tumour, potentially driving them toward dysfunctional differentiation states.

## Discussion

GC signalling in T cells is context and dose-dependent, with endogenous GCs playing both immunoenhancing and immunosuppressive roles^28^. Under physiological conditions, circadian GC elevations promote CD8 T cell trafficking, survival, and activation^29^, while perinatal exposure programs long-term CD8 T cell immunity^30^. However, sustained or excessive GC exposure—as occurs via local extra-adrenal steroid biosynthesis within tumour microenvironments^5^—drives the pathological CD8 T cell dysfunction we describe here.

In this study, we reveal that physiological cortisol signalling fundamentally remodels the transcriptional and chromatin landscapes of human CD8 T cells through a previously unrecognised, ligand-dependent cooperation between the GR and RUNX transcription factors. By integrating RNA sequencing, ChIP sequencing, genetic perturbation, biochemical validation, and single-cell transcriptomic analyses of tumour-infiltrating lymphocytes, we identify a novel GR-RUNX regulatory axis that significantly modulates CD8 T cell function within the tumour microenvironment.

Our findings extend the current understanding of GR biology, demonstrating that GR can regulate gene expression via non-canonical interactions with RUNX transcription factors, rather than exclusively through classical GREs. Traditionally, GR-mediated transcriptional regulation has been attributed to direct DNA binding at GREs or through tethered interactions with factors such as NF-κB and AP-1^1,11^. Here, we provide evidence that, in cortisol-treated human CD8 T cells, GR preferentially occupies genomic regions enriched for RUNX motifs. Co-immunoprecipitation experiments confirm a direct, ligand-dependent interaction between GR and RUNX3, positioning RUNX3 as a critical non-canonical partner that enables GR to access distinct regulatory regions beyond those governed by classical GRE-driven mechanisms. We observed cortisol promotes the interaction between GR and RUNX3. We also observed that AF1 region of NR3C1 and C-terminal TAD/ID region of RUNX3 are required for efficient complex formation. Notably, GR disruption did not significantly alter a subset of cortisol-responsive genes, indicating that cortisol can reshape CD8 T cell transcription through GR-independent mechanism. This is consistent with cortisol may act as a broad cellular stimulus that can elicit rapid non-genomic signalling (second-messenger pathway) and activate downstream transcription factors, or CD8 T cell may use alternate cortisol-sensing and metabolic pathways under long-term cortisol exposure. However, this GR-independent component was distinct from GR-RUNX coregulated module.

RUNX transcription factors are established regulators of immune cell differentiation and function^19^. Our data uncover an unexpected role for RUNX3 in human CD8 T cells as a pivotal mediator of cortisol-induced immunomodulation. Genetic ablation of GR expression abolishes cortisol-induced gene expression changes, confirming direct GR dependency. Moreover, cortisol-induced genes with proximal GR-RUNX motif occupancy include key immunoregulatory factors such as *IL7R, FKBP5, TSC22D3, PIK3IP1* and *AREG*, implicating RUNX as a crucial determinant of GC-driven immunomodulation.

A particularly intriguing observation from our study is the paradoxical regulation of signalling pathways by cortisol in CD8 T cells. Cortisol robustly suppressed inflammatory pathways (e.g., TNF-α/NF-κB and IL-2/STAT5 signalling) while simultaneously activating pathways associated with cell survival and proliferation (mTORC1 signalling, G2/M checkpoint). This dual regulatory pattern suggests that cortisol enforces a metabolic checkpoint state, limiting inflammatory effector functions while maintaining viability and proliferative capacity. Such metabolic reprogramming may facilitate prolonged persistence of dysfunctional or exhausted T cells within tumours rather than productive effector differentiation, thus contributing to impaired antitumour immunity. An important question for future work is whether GR-RUNX co-operation preferentially supports maintenance of dysfunctional states, entry into exhaustion trajectories or both and how this is influenced by antigen load, co-stimulation and cytokines in the tumour microenvironment.

Our findings reveal that RUNX-GR dependent gene regulation is particularly prominent within the tumour microenvironment across multiple cancer types. This tumour-specific upregulation suggests that local factors within the tumour might be driving this regulatory programme. Local steroid biosynthesis within the tumour microenvironment^5,6,9^ provides sustained cortisol exposure inducing the GC signalling pathway, which subsequently activates RUNX-GR-mediated transcriptional programs that influence the fate and function of tumour-infiltrating CD8 T cells. The co-regulation by RUNX factors and GR in tumour-infiltrating T cells is particularly intriguing given that both transcription factor families have been implicated in immune cell differentiation and function. RUNX family transcription factors are known to function redundantly, with RUNX1, RUNX2, and RUNX3 often binding to the same genomic sites and collaboratively regulating similar target genes. GR collaborates with pioneer factors and other transcription factors to regulate target gene expression^31–34^. Here we identify RUNX3 as a critical GR co-factor to regulate cell type-specific transcriptional programs. The observed correlation between RUNX-GR co-regulated gene expression and T cell exhaustion markers suggests that this transcriptional program may contribute to the development of the exhausted phenotype that characterises many tumour-infiltrating T cells. In tumour-infiltrating CD8 T cell subsets we observed GR–RUNX target-gene expressing cells were transcriptionally aligned with pre-dysfunctional state and were distinct from cytotoxic CD8 T cell cluster. Conceptually, this observation supports a model in which cortisol signalling through GR-RUNX modulation reinforces the establishment of an intermediate pre-dysfunctional state that can lead to an exhausted phenotype. However detailed molecular mechanism needs further exploration.

This finding provides new insights into the molecular mechanisms that may underlie T cell dysfunction within the tumour microenvironment and identifies potential therapeutic targets for enhancing anti-tumour immunity. Genetic disruption of RUNX3 in peripheral or tumour-infiltrating CD8 T cell and studying its effects in anti-tumour cytotoxicity in cortisol rich microenvironment will elucidate the mechanism of GR-RUNX mediated CD8 T cell dysfunction. Indeed, recent studies have highlighted local GC production within tumours as an immunosuppressive mechanism exploited by cancer cells to evade immune surveillance^2,5–8,35–39^. Our results provide mechanistic insights into how this local steroid signalling axis may operate at the transcriptional level through GR-RUNX cooperation leading T cell dysfunction.

From a translational perspective, our findings suggest novel therapeutic opportunities aimed at improving cancer immunotherapy efficacy by targeting context-specific GR interactions. Systemic inhibition of GC signalling is challenging as GC maintains physiological homeostasis; thus, selective disruption of GR-RUNX3 interactions or downstream targets identified herein may represent promising therapeutic strategies to enhance antitumour immunity without broadly compromising physiological GC functions. Future studies should explore small-molecule inhibitors or targeted genetic approaches to specifically disrupt GR-RUNX3 interactions in preclinical tumour models. Exploring downstream effectors of RUNX-GR module can reveal new druggable targets to enhance anti-tumour immunity.

In conclusion, our study identifies RUNX3 as a novel non-canonical partner of GR in human CD8 T cells and reveals a previously unrecognised mechanism by which endogenous GCs modulate antitumour immunity. Targeting context-specific transcription factor interactions represents an exciting new avenue for therapeutic intervention aimed at reversing steroid-driven immunosuppression within tumours.

### Limitations of the study

Nevertheless, several questions remain unanswered: First, although we demonstrated robust ligand-dependent GR-RUNX3 interaction *in vitro* and identified direct target genes through integrative genomic analyses, further studies are required to clarify the precise molecular determinants mediating this interaction at endogenous loci and its functional outcome changing chromatin structure at the endogenous locus. Second, additional research is needed to delineate the relative contributions of RUNX family members in mediating GC responses in distinct immunological contexts. Finally, future work should validate our findings using *in vivo* tumour models to assess whether disrupting GR-RUNX interactions can effectively restore CD8 T cell functionality and enhance antitumour responses.

## Methods

### Human CD8 T cell culture

Peripheral blood samples from healthy male adult donors (n=3) were obtained by informed consent. Peripheral blood mononuclear cells (PBMCs) were recovered through density gradient centrifugation using Ficoll-Paque Plus (GE Healthcare). CD8 T cells were isolated from PBMCs. CD8 T cells were isolated from PBMCs by immunomagnetic negative selection (Stem Cell Technologies). CD8 T cells were cultured for a total of 8 days in Immunocult-XF T Cell Expansion media (Stem Cell Technologies) with 50 µg/mL of Gentamicin (Gibco). Isolated cells were stimulated with plate bound anti-CD3 (Biolegend, Clone: OKT3, 1 µg/mL) and anti-CD28 (Biolegend, Clone: CD28.2, 1 µg/mL) for 72 hours. Next, CD8 T Cells were rested and expanded in fresh Immunocult media in the presence of 100 U/mL of human IL-2 (Peprotech) for a further 72 hours. Cells were then re-stimulated with plate bound anti-CD3 (Biolegend, Clone: OKT3 1 µg/mL) and anti-CD28 (Biolegend Clone: CD28.2 1 µg/mL) in the presence of 100 nM of cortisol [hydrocortisone (HC) – Sigma] or vehicle control (methanol, Sigma) for 48 hours. All procedures were approved by the Human Biology Research Ethics Committee of the University of Cambridge (HBREC2019.15).

### CRISPR-Cas9 mediated *NR3C1* deletion

Seventy-two hours after T cell activation, cells were washed with PBS and resuspended in complete P3 Primary Cell Nucleofection Solution (Lonza) according to the manufacturer’s instructions. For each nucleofection, 1 × 10⁶ cells were used. Two distinct ribonucleoprotein (RNP) complexes were prepared, each targeting a specific sgRNA designed for the deletion of NR3C1. The sequences for the sgRNAs were as follows:

1. sgRNA 1: 5’ – CUUUAAGUCUGUUUCCCCCG – 3’
2. sgRNA 2: 5’– AACCUCCAACAGUGACACCA – 3’

Both sgRNAs were synthesised by Synthego, and the Alt-R S.p. High Fidelity Cas9 Nuclease V3 (Integrated DNA Technologies) was used to form the RNP complexes. Each RNP complex was assembled at a 3:1 molar ratio of sgRNA to Cas9 (2.48 µM Cas9 + 7.52 µM sgRNA). The prepared Cas9 RNP complexes were mixed with the cells and transferred to a 20 µL Nucleocuvette Vessel. Electroporation was performed using a 4D Nucleofector with the EO-115 program (Lonza). For control experiments, NR3C1-sufficient CD8 T cells were processed in parallel and electroporated with High Fidelity Cas9 protein only (Mock Control). Immediately following nucleofection, 80 µL of pre-warmed ImmunoCult-XF media (without supplements) was added directly to the Nucleocuvette Vessel, and the cells were allowed to recover at 37°C for 20 minutes. The cells were then transferred to a 24-well plate and cultured for 72 hours in fresh ImmunoCult-XF media supplemented with 100 U/mL of IL-2 (PeproTech) and 50 µg/mL of Gentamicin (Gibco™).

### Single-cell RNA sequencing (scRNA-seq), clustering and visualisation

The previously published and publicly available scRNA-seq datasets used in this study were as follows GSE164690^40–42^, GSE161529^43,44^, GSE127465^45^, GSE155698^46,47^ and an integrated scRNA seq of NSCLC from 103 patients^48^. The data has been downloaded from NCBI GEO (https://www.ncbi.nlm.nih.gov/geo/) and processed with the standard scRNA-seq integration pipeline in Seurat. To cluster the cells, we first identified the top 2,000 most variable genes using the FindVariableFeatures function in Seurat. For cell visualization, we applied the Uniform Manifold Approximation and Projection (UMAP) algorithm to project the cells’ PCA space representation into two dimensions. Cell clustering in PCA space was performed with the Shared Nearest Neighbor (SNN) algorithm, using Seurat V3’s FindNeighbors and FindClusters functions. We then visualised the resulting clusters in UMAP space with the DimPlot function. The differentially expressed genes in each cluster were identified with the FindAllMarkers function in Seurat. The clusters have been identified either by previous annotation or based on the Cluster Identity Predictor (CIPR) program and has been validated based on marker gene expression from Panglo DB. Gene expression levels were visualised using heatmaps, violin plots, and bar plots, created with the pheatmap package (version 1.0.12) (available at https://cran.r-project.org/web/packages/pheatmap) and the dittoSeq R package (version 1.2.4). Data scaling during visualization was performed automatically using the default settings of these packages. Cell type information was retrieved from the metadata slot of the downloaded object. To quantify GR-RUNX3 co-expression within tumour-infiltrating CD8 T Cells, we calculated the co-expression score as the geometric mean of log-normalised counts in single cell expression of *NR3C1* and *RUNX3*. Gene module scores for subpopulations, as defined above, were calculated using the Seurat function AddModuleScore. CD8 T-cell transcriptional states were quantified in a Seurat object using gene-signature module scoring.

Marker gene sets for CD8 T-cell states

Curated marker genes were compiled a priori to represent major CD8 T-cell states. The “general” marker sets comprised:

**Naïve-like:** *TCF7, CCR7, SELL, LEF1, IL7R, LTB*

**Pre-dysfunctional:** *GZMK, ZNF683, CD28, FYN, EOMES, CXCR3, CXCR4, CD44*

**Dysfunctional/exhausted:** *PDCD1, HAVCR2, LAG3, TIGIT, CTLA4, LAYN, CXCL13, ENTPD1, ITGAE, IFNG*

**Cytotoxic:** *CX3CR1, PRF1, GZMA, GZMB, GZMH, GNLY, FGFBP2, NKG7, KLRD1, FCGR3A, KLRG1*

Marker-gene expression and module scores were visualised on UMAP embeddings using FeaturePlot.

### Single-cell trajectory analysis

Single-cell trajectory analysis was performed using Monocle (version 2.20.0). Briefly, size factors and dispersions were estimated with default settings. Differentially expressed genes were identified using Monocle’s differentialGeneTest function (q-value < 0.01). Identified genes were used as ordering genes. Dimensionality was reduced using the DDRTree algorithm, and trajectories were constructed by ordering cells based on pseudotime.

### Western blot analysis

72 hours post nucleofection, CD8 T cells were washed once with ice cold PBS and lysed in radioimmunoprecipitation assay (RIPA) buffer (Merck) with proteinase and phosphatase inhibitors (Roche). Protein concentrations were determined by Pierce™ BCA Protein Assay Kit and performed according to the manufacturer’s instructions (Thermo-Scientific). Protein samples (15 µg) were separated on a NuPAGE 4-12% Bis-Tris polyacrylamide gels (Invitrogen), transferred onto nitrocellulose membranes (Bio-Rad) and probed using rabbit anti-GR (Clone: D6H2L, Cell Signaling Technology) and mouse beta-actin (Clone: 8H10, Origene) antibodies. Detection of immunoreactive bands was performed using secondary goat anti-rabbit IRDye™ 680CW and goat anti-mouse IRDye™ 800CW antibodies. Images acquired on the Odyssey DLx imaging system (Li-COR).

### Bulk RNA sequencing analysis

Cell pellets were layered with 40 µL of RNAlater Stabilization solution (Invitrogen) and stored at -80°C prior to RNA extraction. Samples were homogenised using the QIAshredder Kit (Qiagen) according to manufacturer’s instructions. Next, total RNA was extracted using the RNeasy Plus Mini Kit (Qiagen). RNA sequencing was performed in paired-end on Illumina NovaSeq 6000 platform, yielding 40 million reads per sample (Novogene, UK). Raw sequencing reads were quality-controlled with FastQC (version 0.11.9), followed by alignment to the reference genome gencode version 42 (Homo sapiens genome hg38/GRCh38) using the alignment tool hisat2 (version 2.2.1). The sam files were further inspected, filtered, and processed using samtools (version 1.13). HTseq counts obtained from bam files were annotated and subjected to DESeq2 (version 1.34.0) analysis in R (version 4.1.1). For DEseq2 analysis, for each gene, a log2 fold change was computed and the significance level was assessed using the Wald test statistic with a p value and an adjusted p value. Differentially expressed genes across different conditions have been identified using the cutoff of a p-adjusted value of 0.05. The volcano plot has been constructed in GraphPad Prism 9.5.1 based on the differentially expressed genes found in bulk RNA sequencing. After differential expression analysis, genes were ranked according to their fold change and by using the fGSEA algorithm, we performed Gene Set Enrichment Analysis (GSEA). The human collection H (hallmark gene sets) from the Molecular Signatures Database (MSigDB) were used to identify positively and negatively enriched pathways. All steps were performed in R version 4.1.1.

### Chromatin immunoprecipitation (ChIP) followed by sequencing

Chromatin immunoprecipitation (ChIP) was performed using the SimpleChIP Enzymatic Chromatin IP Kit (Cell Signaling Technology) following the manufacturer’s protocol with minor modifications. Briefly, cells were crosslinked with 1% formaldehyde (Merck) in serum-free RPMI-1640 for 10 min at room temperature (RT) and quenched with 1X glycine for 5 min. Nuclei were isolated and chromatin digested with 0.3 µL micrococcal nuclease in 200 µL ChIP Buffer at 37 °C for 20 min, followed by brief sonication (Branson 450-D sonicator, amplitude 36%, 4 × 5 s pulses on ice). Chromatin was immunoprecipitated overnight at 4 °C with 1 µg each of anti-GR antibodies (PA1-511A, Thermo-Scientific; D6H2L, Cell Signaling Technology). Immune complexes were captured with ChIP-Grade Protein G Magnetic Beads (2 h, 4 °C), washed sequentially with low-salt and high-salt buffers, and eluted with 1X ChIP Elution Buffer at 65 °C for 30 min. Crosslinks were reversed at 65 °C for 2 hours (5 M NaCl, Proteinase K), with DNA purified and submitted for Illumina sequencing (Novogene, Cambridge, UK).

### ChIP-seq data analysis

Raw reads from fastq files were first trimmed using fastp 0.20.0 with the setting ‘-l 25 --detect_adapter_for_pe’. Trimmed reads were mapped to the human genome hg38 by hisat2 2.1.0 with default setting and sorted by samtools 1.9 with flag ‘-ShuF 4 -q 30 -f 2’. After mapping, duplicated reads were removed by the picard tool (v.2.20.8) (http://broadinstitute.github.io/picard/). Peaks were then called using MACS2 and annotated using HOMER^49^.

### Integration of bulk RNA-seq and ChIP-seq data

After analysing bulk RNA-Seq and ChIP-Seq data, we compared the differentially expressed genes in RNA-seq of cortisol treated vs. vehicle control and all annotated peaks in ChIP-seq data. Downstream analysis was performed based on the genes between the RNA-seq and HC-treated ChIP-seq peaks, which include a total of 35 genes. The venn diagram to visualise the common genes was constructed using InteractiVenn. A heatmap has been drawn in gplots (version 3.1.3) in R version 4.1.1 to show the normalised counts of genes (transformed into row z-score) common in differentially expressed genes in RNA-seq and annotated peaks from cortisol (HC) treated cells. The protein-protein interaction (PPI) network has been constructed in STRING (version 11.5) and visualised in cytoscape (version 3.91). All 35 differentially expressed genes have been used as inputs in STRING, but only 8 genes constitute an intact network. The pie chart demonstrating the genomic position diversity of peaks obtained from cortisol treated cells in ChIP-seq data has been constructed in GraphPad Prism 9.5.1.

### Overexpression of NR3C1 and RUNX3 in HEK-293T

A custom polycistronic plasmid (EF1A > hRUNX3 [NM_001320672.1] : IRES : hNR3C1 [NM_001364185.1] : IRES : EGFP) was designed and generated by VectorBuilder. Driven by the EF1A promoter, this construct produces a single mRNA transcript translated into RUNX3, NR3C1, and EGFP via IRES elements. HEK-293T cells were transfected with 5 μg of plasmid using Lipofectamine 3000 (Invitrogen) in serum-free Opti-MEM, following the manufacturer’s protocol. Transfection was performed at 70–80% confluence. After 6 hours, 100nM of cortisol (Sigma-Aldrich) was added directly to the culture medium, and cells were incubated for an additional 48 hours to allow for cortisol-mediated effects.

### Co-Immunoprecipitation (Co-IP)

Cells were harvested by washing with ice-cold PBS and collected by centrifugation at 300 × g for 5 minutes at 4 °C. Cell lysis was performed using Pierce IP Lysis Buffer (Thermo Scientific) supplemented with protease and PhosSTOP phosphatase inhibitors (Roche). Lysates were incubated on ice for 30 minutes, followed by mild sonication (three 5 s pulses at 36% amplitude) using a Branson 4D Sonicator. Lysates were cleared by centrifugation at 16,000 × g for 20 minutes at 4 °C. The supernatant was pre-cleared by incubation with 10 μL of Protein A/G Plus-Agarose Beads (Santa Cruz) for 1 hour at 4 °C with rotation, followed by centrifugation at 3,000 × g for 10 minutes. Clarified lysates were incubated overnight at 4 °C with either anti-NR3C1 (D6H2L, Cell Signaling Technology; 1:100) or control rabbit IgG (Cell Signaling Technology), followed by addition of 30 μL Protein A/G Plus-Agarose Beads and further incubation for 2 hours at 4 °C. Beads were washed three times with lysis buffer, and bound proteins were eluted in Laemmli sample buffer. Eluates were analysed by western blotting using anti-RUNX1/AML1, RUNX3, and RUNX2 antibodies (EPR3099, Abcam). The details of the mutations (ΔAF1-NR3C1, ΔTAD-RUNX3, ΔTAD ID-RUNX3) has been provided in the supplementary document.

## Resource availability

### Lead contact

Further information and requests for resources and reagents should be directed to and will be fulfilled by the Lead Contact, Bidesh Mahata

## Material availability

This study did not generate unique reagents.

## Data and code availability

◦ Data: Newly generated bulk RNA seq and ChIP seq data are deposited to EBI ArrayExpress with the accession numbers E-MTAB-16594, E-MTAB-16635 (https://www.ebi.ac.uk/biostudies/arrayexpress).
◦ Code: This paper does not report original code.
◦ Additional information: Any additional information required to reanalyse the data reported in this paper is available from the lead contact upon request.

## Author contribution

CJW, SC, SKS, AAD, CVV, QZ, JP, XC and BM performed the experiments, analysed, interpreted & visualised the data. BM conceptualised and supervised the study. All authors read and approved the draft manuscript before submission.

## Acknowledgements

We would like to thank Dr Andrew Conway-Morris, University of Cambridge for providing ethical support; Amit Mandal, University of Oxford, for critical suggestion and advice on RNA-seq data analysis. We used BioRender.com to generate the graphical illustrations presented in this manuscript.

## Data and materials availability

All newly generated data such as RNA sequencing and ChIP sequencing data used in the study will be available in the public database during revised submission stage.

## Supplementary tables

Supplementary Table S1: Cortisol-induced genes in human CD8 T cell

Supplementary Table S2: Cortisol-induced hallmark pathways in CD8 T cell

Supplementary Table S3: Glucocorticoid binding peaks in CD8 T cell

Supplementary Table S4: Glucocorticoid direct target genes in CD8 T cell

Supplementary Table S5: Cortisol induced genes, ChIP-seq peaks, RUNX regulated genes

---

**Supplementary Figure S1.**
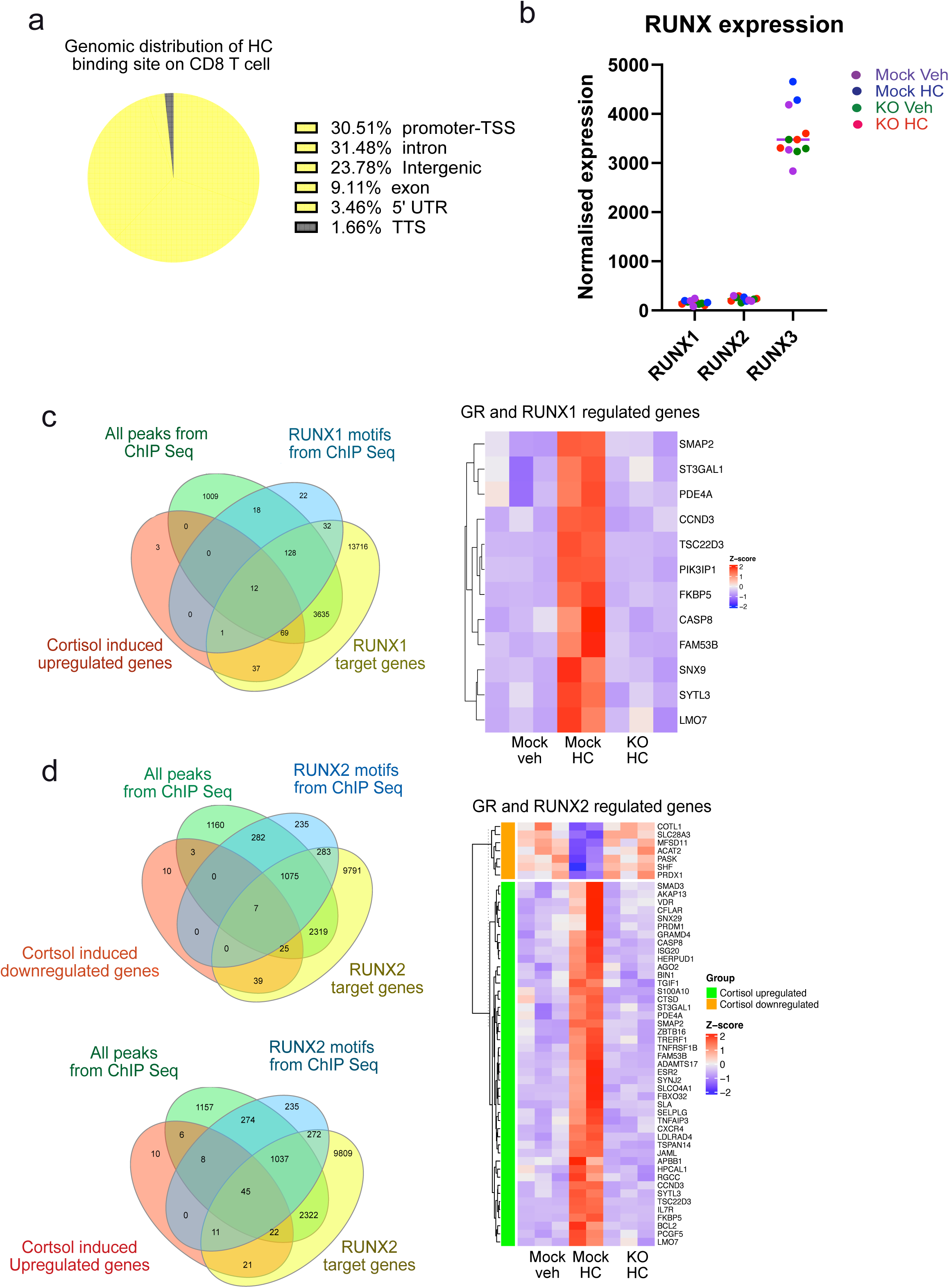
Comparative *RUNX 1, 2* and *3* expression and identification of GR-RUNX 1 and 2 co-regulated genes. **a.** Genomic distribution of GR binding sites in HC-treated CD8 T cells. **b.** Comparative *RUNX 1, 2* and *3* expression in activated CD8 T cells in the RNAseq. Normalised expression of these genes from different donors has been plotted here **c.** Venn diagram depicting the set of genes jointly regulated by the GR and RUNX1, identified through an integrative analysis of cortisol-induced upregulated genes, GR ChIP-seq peaks, GR peaks containing RUNX1 motifs, and known RUNX1 target genes. Expression levels of the overlapping gene set are visualised in a heatmap across multiple experimental conditions. **d.** Venn diagram illustrating the genes co-regulated by GR and RUNX2, identified by integrating cortisol-responsive genes (both upregulated and downregulated), GR ChIP-seq data, GR peaks with RUNX2 motifs, and annotated RUNX2 targets. Expression patterns of these shared genes under different conditions are shown in the accompanying heatmap. Specification of the samples: KO denotes glucocorticoid receptor (GR) knockout, HC denotes hydrocortisone (cortisol) treated. Mock denotes wildtype. Veh denotes vehicle.

**Supplementary Figure S2.**
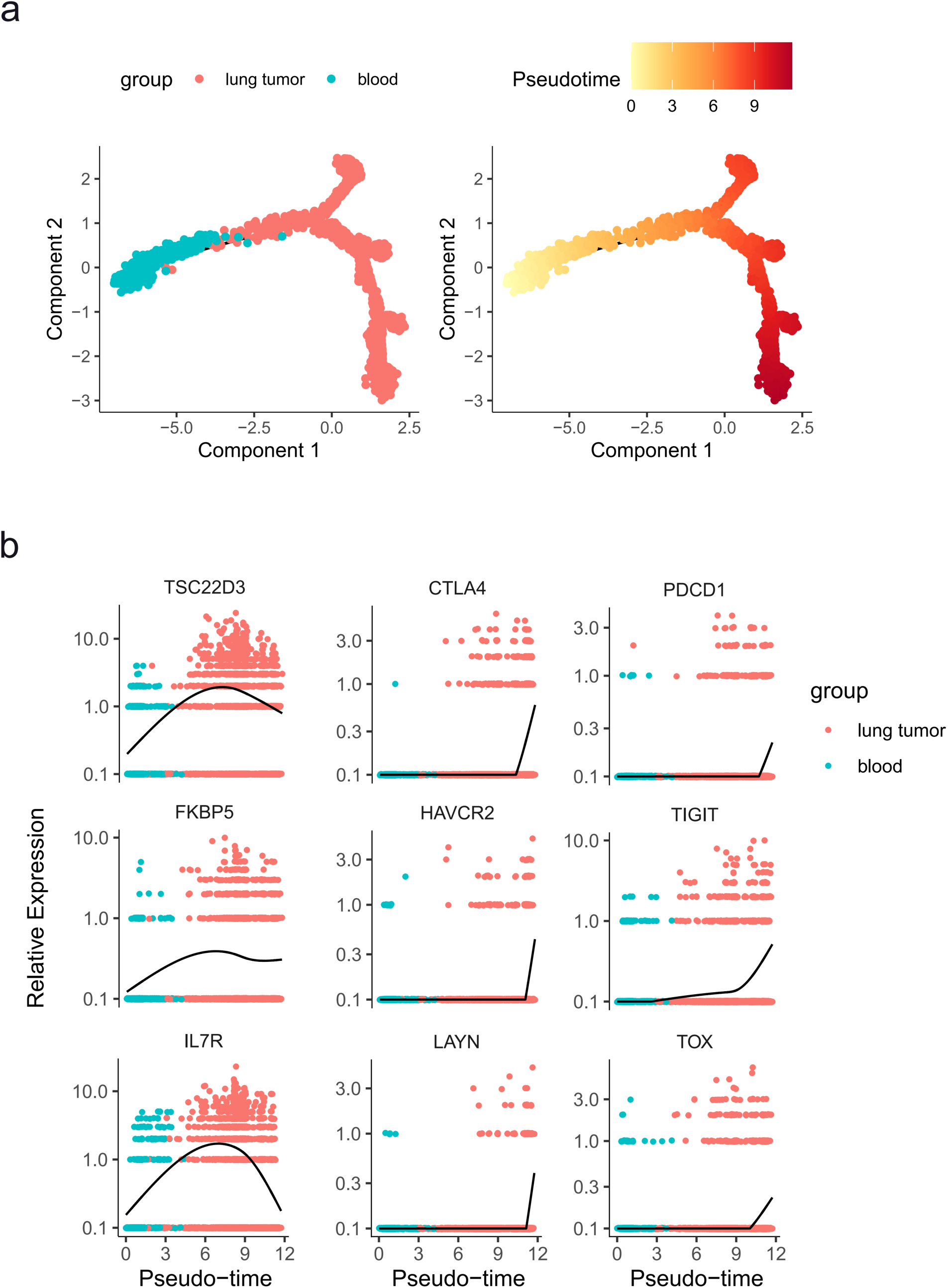
Single-cell trajectory analysis of CD8 T cells from lung tumours and blood samples. **a.** Pseudo-temporal trajectories of CD8 T cells inferred using Monocle, coloured by sample origin and pseudotime. **b.** Expression patterns of cortisol-responsive genes (*TSC22D3, IL7R, FKBP5*) and inhibitory receptors (*PDCD1, CTLA4, HAVCR2, TIGIT, LAYN, TOX*) along the inferred trajectories.

**Supplementary Figure S3.**
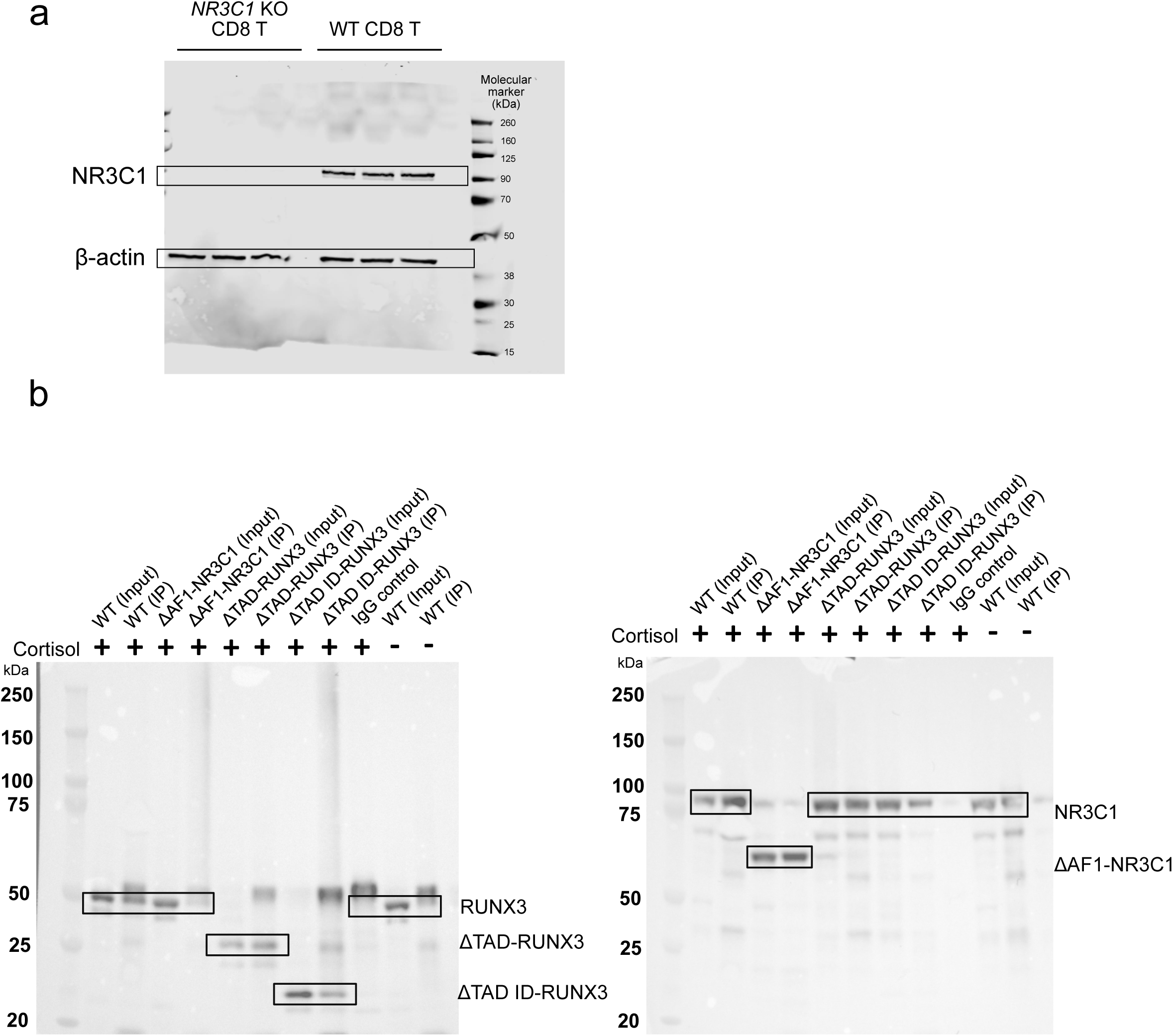
Original western blots. **a.** This western blot corresponds to Figure 1d. **b.** These western blots correspond to Figure 3c.

## References

1. Cain, D.W., and Cidlowski, J.A. (2017). Immune regulation by glucocorticoids. Nat Rev Immunol 17, 233–247. 10.1038/nri.2017.1.

2. Vandewalle, J., Luypaert, A., De Bosscher, K., and Libert, C. (2018). Therapeutic Mechanisms of Glucocorticoids. Trends Endocrinol Metab 29, 42–54. 10.1016/j.tem.2017.10.010.

3. Quatrini, L., and Ugolini, S. (2021). New insights into the cell- and tissue-specificity of glucocorticoid actions. Cell Mol Immunol 18, 269–278. 10.1038/s41423-020-00526-2.

4. Obradovic, M.M.S., Hamelin, B., Manevski, N., Couto, J.P., Sethi, A., Coissieux, M.M., Munst, S., Okamoto, R., Kohler, H., Schmidt, A., and Bentires-Alj, M. (2019). Glucocorticoids promote breast cancer metastasis. Nature 567, 540–544. 10.1038/s41586-019-1019-4.

5. Acharya, N., Madi, A., Zhang, H., Klapholz, M., Escobar, G., Dulberg, S., Christian, E., Ferreira, M., Dixon, K.O., Fell, G., et al. (2020). Endogenous Glucocorticoid Signaling Regulates CD8(+) T Cell Differentiation and Development of Dysfunction in the Tumor Microenvironment. Immunity 53, 658–671 e656. 10.1016/j.immuni.2020.08.005.

6. Mahata, B., Pramanik, J., van der Weyden, L., Polanski, K., Kar, G., Riedel, A., Chen, X., Fonseca, N.A., Kundu, K., Campos, L.S., et al. (2020). Tumors induce de novo steroid biosynthesis in T cells to evade immunity. Nat Commun 11, 3588. 10.1038/s41467-020-17339-6.

7. Chakraborty, S., Pramanik, J., and Mahata, B. (2021). Revisiting steroidogenesis and its role in immune regulation with the advanced tools and technologies. Genes Immun 22, 125–140. 10.1038/s41435-021-00139-3.

8. Pramanik, J., Shaji, S.K., Zaman, M., Brown, B., Zhang, B., Yamashita-Kanemaru, Y., Homer, N.Z.M., Hussein, H.A.M., Zhao, Q., Okkenhaug, K., et al. (2025). Drug repurposing reveals posaconazole as a CYP11A1 inhibitor enhancing anti-tumor immunity. iScience 28, 112488. 10.1016/j.isci.2025.112488.

9. Zhao, Q., Pramanik, J., Lu, Y., Homer, N.Z.M., Imianowski, C.J., Zhang, B., Iqbal, M., Shaji, S.K., Morris, A.C., Roychoudhuri, R., et al. (2025). Perturbing local steroidogenesis to improve breast cancer immunity. Nat Commun 16, 3945. 10.1038/s41467-025-59356-3.

10. Sandor, L.F., Huh, J.B., Benko, P., Hiraga, T., Poliska, S., Dobo-Nagy, C., Simpson, J.P., Homer, N.Z.M., Mahata, B., and Gyori, D.S. (2024). De novo steroidogenesis in tumor cells drives bone metastasis and osteoclastogenesis. Cell Rep 43, 113936. 10.1016/j.celrep.2024.113936.

11. Weikum, E.R., Knuesel, M.T., Ortlund, E.A., and Yamamoto, K.R. (2017). Glucocorticoid receptor control of transcription: precision and plasticity via allostery. Nat Rev Mol Cell Biol 18, 159–174. 10.1038/nrm.2016.152.

12. De Bosscher, K., Beck, I.M., Dejager, L., Bougarne, N., Gaigneaux, A., Chateauvieux, S., Ratman, D., Bracke, M., Tavernier, J., Vanden Berghe, W., et al. (2014). Selective modulation of the glucocorticoid receptor can distinguish between transrepression of NF-kappaB and AP-1. Cell Mol Life Sci 71, 143–163. 10.1007/s00018-013-1367-4.

13. Escoter-Torres, L., Greulich, F., Quagliarini, F., Wierer, M., and Uhlenhaut, N.H. (2020). Anti-inflammatory functions of the glucocorticoid receptor require DNA binding. Nucleic Acids Res 48, 8393–8407. 10.1093/nar/gkaa565.

14. Franco, L.M., Gadkari, M., Howe, K.N., Sun, J., Kardava, L., Kumar, P., Kumari, S., Hu, Z., Fraser, I.D.C., Moir, S., et al. (2019). Immune regulation by glucocorticoids can be linked to cell type-dependent transcriptional responses. J Exp Med 216, 384–406. 10.1084/jem.20180595.

15. Wherry, E.J., and Kurachi, M. (2015). Molecular and cellular insights into T cell exhaustion. Nat Rev Immunol 15, 486–499. 10.1038/nri3862.

16. Philip, M., and Schietinger, A. (2022). CD8(+) T cell differentiation and dysfunction in cancer. Nat Rev Immunol 22, 209–223. 10.1038/s41577-021-00574-3.

17. Giles, A.J., Hutchinson, M.N.D., Sonnemann, H.M., Jung, J., Fecci, P.E., Ratnam, N.M., Zhang, W., Song, H., Bailey, R., Davis, D., et al. (2018). Dexamethasone-induced immunosuppression: mechanisms and implications for immunotherapy. J Immunother Cancer 6, 51. 10.1186/s40425-018-0371-5.

18. Xia, M., Gasser, J., and Feige, U. (1999). Dexamethasone enhances CTLA-4 expression during T cell activation. Cell Mol Life Sci 55, 1649–1656. 10.1007/s000180050403.

19. Seo, W., and Taniuchi, I. (2020). The Roles of RUNX Family Proteins in Development of Immune Cells. Mol Cells 43, 107–113. 10.14348/molcells.2019.0291.

20. Mevel, R., Draper, J.E., Lie, A.L.M., Kouskoff, V., and Lacaud, G. (2019). RUNX transcription factors: orchestrators of development. Development 146. 10.1242/dev.148296.

21. Korinfskaya, S., Parameswaran, S., Weirauch, M.T., and Barski, A. (2021). Runx Transcription Factors in T Cells-What Is Beyond Thymic Development? Front Immunol 12, 701924. 10.3389/fimmu.2021.701924.

22. Woolf, E., Xiao, C., Fainaru, O., Lotem, J., Rosen, D., Negreanu, V., Bernstein, Y., Goldenberg, D., Brenner, O., Berke, G., et al. (2003). Runx3 and Runx1 are required for CD8 T cell development during thymopoiesis. Proc Natl Acad Sci U S A 100, 7731–7736. 10.1073/pnas.1232420100.

23. Milner, J.J., Toma, C., Yu, B., Zhang, K., Omilusik, K., Phan, A.T., Wang, D., Getzler, A.J., Nguyen, T., Crotty, S., et al. (2017). Runx3 programs CD8(+) T cell residency in non-lymphoid tissues and tumours. Nature 552, 253–257. 10.1038/nature24993.

24. Egawa, T., Tillman, R.E., Naoe, Y., Taniuchi, I., and Littman, D.R. (2007). The role of the Runx transcription factors in thymocyte differentiation and in homeostasis of naive T cells. J Exp Med 204, 1945–1957. 10.1084/jem.20070133.

25. Lu, L., Wen, Y., Yao, Y., Chen, F., Wang, G., Wu, F., Wu, J., Narayanan, P., Redell, M., Mo, Q., and Song, Y. (2018). Glucocorticoids Inhibit Oncogenic RUNX1-ETO in Acute Myeloid Leukemia with Chromosome Translocation t(8;21). Theranostics 8, 2189–2201. 10.7150/thno.22800.

26. Khan, S.H., Ling, J., and Kumar, R. (2011). TBP binding-induced folding of the glucocorticoid receptor AF1 domain facilitates its interaction with steroid receptor coactivator-1. PLoS One 6, e21939. 10.1371/journal.pone.0021939.

27. van der Leun, A.M., Thommen, D.S., and Schumacher, T.N. (2020). CD8(+) T cell states in human cancer: insights from single-cell analysis. Nat Rev Cancer 20, 218–232. 10.1038/s41568-019-0235-4.

28. Taves, M.D., and Ashwell, J.D. (2021). Glucocorticoids in T cell development, differentiation and function. Nat Rev Immunol 21, 233–243. 10.1038/s41577-020-00464-0.

29. Shimba, A., Cui, G., Tani-Ichi, S., Ogawa, M., Abe, S., Okazaki, F., Kitano, S., Miyachi, H., Yamada, H., Hara, T., et al. (2018). Glucocorticoids Drive Diurnal Oscillations in T Cell Distribution and Responses by Inducing Interleukin-7 Receptor and CXCR4. Immunity 48, 286–298 e286. 10.1016/j.immuni.2018.01.004.

30. Hong, J.Y., Lim, J., Carvalho, F., Cho, J.Y., Vaidyanathan, B., Yu, S., Annicelli, C., Ip, W.K.E., and Medzhitov, R. (2020). Long-Term Programming of CD8 T Cell Immunity by Perinatal Exposure to Glucocorticoids. Cell 180, 847–861 e815. 10.1016/j.cell.2020.02.018.

31. John, S., Sabo, P.J., Thurman, R.E., Sung, M.H., Biddie, S.C., Johnson, T.A., Hager, G.L., and Stamatoyannopoulos, J.A. (2011). Chromatin accessibility pre-determines glucocorticoid receptor binding patterns. Nat Genet 43, 264–268. 10.1038/ng.759.

32. Stavreva, D.A., Coulon, A., Baek, S., Sung, M.H., John, S., Stixova, L., Tesikova, M., Hakim, O., Miranda, T., Hawkins, M., et al. (2015). Dynamics of chromatin accessibility and long-range interactions in response to glucocorticoid pulsing. Genome Res 25, 845–857. 10.1101/gr.184168.114.

33. Voss, T.C., and Hager, G.L. (2014). Dynamic regulation of transcriptional states by chromatin and transcription factors. Nat Rev Genet 15, 69–81. 10.1038/nrg3623.

34. Biddie, S.C., John, S., Sabo, P.J., Thurman, R.E., Johnson, T.A., Schiltz, R.L., Miranda, T.B., Sung, M.H., Trump, S., Lightman, S.L., et al. (2011). Transcription factor AP1 potentiates chromatin accessibility and glucocorticoid receptor binding. Mol Cell 43, 145–155. 10.1016/j.molcel.2011.06.016.

35. Ahmed, A., Reinhold, C., Breunig, E., Phan, T.S., Dietrich, L., Kostadinova, F., Urwyler, C., Merk, V.M., Noti, M., Toja da Silva, I., et al. (2023). Immune escape of colorectal tumours via local LRH-1/Cyp11b1-mediated synthesis of immunosuppressive glucocorticoids. Mol Oncol 17, 1545–1566. 10.1002/1878-0261.13414.

36. Michalek, S., Goj, T., Plazzo, A.P., Marovca, B., Bornhauser, B., and Brunner, T. (2022). LRH-1/NR5A2 interacts with the glucocorticoid receptor to regulate glucocorticoid resistance. EMBO Rep 23, e54195. 10.15252/embr.202154195.

37. Taves, M.D., Otsuka, S., Taylor, M.A., Donahue, K.M., Meyer, T.J., Cam, M.C., and Ashwell, J.D. (2023). Tumors produce glucocorticoids by metabolite recycling, not synthesis, and activate Tregs to promote growth. J Clin Invest 133. 10.1172/JCI164599.

38. Merk, V.M., Grob, L., Fleischmann, A., and Brunner, T. (2023). Human lung carcinomas synthesize immunoregulatory glucocorticoids. Genes Immun 24, 52–56. 10.1038/s41435-023-00194-y.

39. Sidler, D., Renzulli, P., Schnoz, C., Berger, B., Schneider-Jakob, S., Fluck, C., Inderbitzin, D., Corazza, N., Candinas, D., and Brunner, T. (2011). Colon cancer cells produce immunoregulatory glucocorticoids. Oncogene 30, 2411–2419. 10.1038/onc.2010.629.

40. Kurten, C.H.L., Kulkarni, A., Cillo, A.R., Santos, P.M., Roble, A.K., Onkar, S., Reeder, C., Lang, S., Chen, X., Duvvuri, U., et al. (2021). Investigating immune and non-immune cell interactions in head and neck tumors by single-cell RNA sequencing. Nat Commun 12, 7338. 10.1038/s41467-021-27619-4.

41. Janjic, B.M., Kulkarni, A., Ferris, R.L., Vujanovic, L., and Vujanovic, N.L. (2022). Human B Cells Mediate Innate Anti-Cancer Cytotoxicity Through Concurrent Engagement of Multiple TNF Superfamily Ligands. Front Immunol 13, 837842. 10.3389/fimmu.2022.837842.

42. Wang, J., Li, H., Kulkarni, A., Anderson, J.L., Upadhyay, P., Onyekachi, O.V., Arantes, L., Banerjee, H., Kane, L.P., Zhang, X., et al. (2025). Differential impact of TIM-3 ligands on NK cell function. J Immunother Cancer 13. 10.1136/jitc-2024-010618.

43. Pal, B., Chen, Y., Vaillant, F., Capaldo, B.D., Joyce, R., Song, X., Bryant, V.L., Penington, J.S., Di Stefano, L., Tubau Ribera, N., et al. (2021). A single-cell RNA expression atlas of normal, preneoplastic and tumorigenic states in the human breast. EMBO J 40, e107333. 10.15252/embj.2020107333.

44. Chen, Y., Pal, B., Lindeman, G.J., Visvader, J.E., and Smyth, G.K. (2022). R code and downstream analysis objects for the scRNA-seq atlas of normal and tumorigenic human breast tissue. Sci Data 9, 96. 10.1038/s41597-022-01236-2.

45. Zilionis, R., Engblom, C., Pfirschke, C., Savova, V., Zemmour, D., Saatcioglu, H.D., Krishnan, I., Maroni, G., Meyerovitz, C.V., Kerwin, C.M., et al. (2019). Single-Cell Transcriptomics of Human and Mouse Lung Cancers Reveals Conserved Myeloid Populations across Individuals and Species. Immunity 50, 1317–1334 e1310. 10.1016/j.immuni.2019.03.009.

46. Steele, N.G., Carpenter, E.S., Kemp, S.B., Sirihorachai, V.R., The, S., Delrosario, L., Lazarus, J., Amir, E.D., Gunchick, V., Espinoza, C., et al. (2020). Multimodal Mapping of the Tumor and Peripheral Blood Immune Landscape in Human Pancreatic Cancer. Nat Cancer 1, 1097–1112. 10.1038/s43018-020-00121-4.

47. Halbrook, C.J., Thurston, G., Boyer, S., Anaraki, C., Jimenez, J.A., McCarthy, A., Steele, N.G., Kerk, S.A., Hong, H.S., Lin, L., et al. (2022). Differential integrated stress response and asparagine production drive symbiosis and therapy resistance of pancreatic adenocarcinoma cells. Nat Cancer 3, 1386–1403. 10.1038/s43018-022-00463-1.

48. Prazanowska, K.H., and Lim, S.B. (2023). An integrated single-cell transcriptomic dataset for non-small cell lung cancer. Sci Data 10, 167. 10.1038/s41597-023-02074-6.

49. Pramanik, J., Chen, X., Kar, G., Henriksson, J., Gomes, T., Park, J.E., Natarajan, K., Meyer, K.B., Miao, Z., McKenzie, A.N.J., et al. (2018). Genome-wide analyses reveal the IRE1a-XBP1 pathway promotes T helper cell differentiation by resolving secretory stress and accelerating proliferation. Genome Med 10, 76. 10.1186/s13073-018-0589-3.

